# Emotional modulation of visual cortical reactivation during true and false memories

**DOI:** 10.64898/2026.01.08.694977

**Authors:** Christoph Hofstetter, Monika Riegel, Kinga Igloi, Aline Pichon, Patrik Vuilleumier

## Abstract

Previous research revealed that memory retrieval is associated with a reactivation of sensory cortices reflecting the content of the memory. Further, such reactivation is stronger when retrieval is cued with emotional information as compared with neutral information. However, past studies focused on correct retrieval, and it remains unknown whether a similar reactivation occurs during memory errors, and whether this is enhanced in emotional contexts. Here, we hypothesized that emotion may influence memory errors due to an increase of sensory cortical reactivation at the cost of recovering episodic details (i.e. leading to incorrect retrieval of non-specific associations shared with other cues).

We compared brain activations during successful memory retrieval and memory errors in 18 healthy participants who encoded unique negative or neutral scenes paired with one of four associated faces, shared with other trials. Participants were tested the next day with a scene-cued associative retrieval test, while undergoing functional magnetic resonance imaging (fMRI). Results from a face-responsive region of interest in fusiform cortex (unlike a control place-responsive region in parahippocampal cortex) showed stronger activation for both Hits (“Old” | Target) and False Alarms (“Old” | Foil), as compared with Misses (“New” | Target) and Correct Rejections (“New” | Foil). The emotional content of scene cues did not modulate activation patterns during correct retrieval but produced specific effects on Misattribution errors (incorrect “Old” | Target, i.e., retrieval of an incorrect face cued by a previously seen scene). Specifically, Misattributions were accompanied by increased fusiform activation comparable to Hits when cued by negative scenes, but low fusiform activation similar to Misses when cued by neutral scenes. Unlike Misattributions, however, Hits cued by emotional information recruited a distinctive network previously linked to personal episodic memories, including medial prefrontal, cingulate, and hippocampal regions, together with enhanced functional coupling between the latter and fusiform cortex. No implicit effect of previous face exposure was observed in the fusiform during Misses, regardless of emotion context.

Our results suggest that the reactivation of sensory areas follows subjective memory experience during retrieval, and that emotion can enhance such mechanism when the cue is recognized but detailed episodic memory mediated by fronto-hippocampal circuits is absent, leading to the retrieval of inappropriate associations and false memories.

## 1 Introduction

Humans have an extraordinary ability to vividly re-enact sensory details of prior experience. The theory of sensory reinstatement (Danker & Anderson, 2010; Tulving & Thomson, 1973) posits that remembering is associated with a reactivation of sensory information perceived during memory encoding, a process thought to result from hippocampally driven reactivation of cortical areas (Bosch et al., 2014; McClelland et al., 1995; Staresina & Wimber, 2019; Xue, 2018). Accordingly, previous neuroimaging studies showed that brain activity patterns from encoding are reactivated during retrieval in a stimulus-specific manner (Bone et al., 2020). This reactivation may reflect the modality of memories, e.g. the retrieval of visual or auditory information differentially recruits visual or auditory areas, as well as specific content of memories, e.g., retrieval of faces vs. words engage distinct visual areas (Chen et al., 2024; Hofstetter et al., 2012; Polyn et al., 2005). However, the functional significance of such reactivations and their relationship with behavioral performance remain unresolved. Content-specific reactivation has been reported for successful retrieval, but it is unknown whether similar reactivation occurs during incorrect retrieval (i.e. during misses, false alarms and misattributions) and, if so, what would be its role in memory experience or specificity.

One possibility is that cortical reactivation occurs even without conscious recollection, in line with evidence for implicit forms of memory (Bower, 1996; Henke et al., 2003; Paller & Voss, 2004). Neural mechanisms of implicit memory were demonstrated by priming tasks, where performance is influenced by previous exposure without conscious retrieval (Schacter & Buckner, 1998), an effect typically attributed to memory traces in cortical areas previously engaged by the same stimulus (Buckner et al., 1998; Vuilleumier et al., 2002). Another possibility is that cortical reactivation does not reflect veridical memory but rather reconstructive processes at play during remembering (Favila et al., 2020; Xue, 2022). In this case, it might occur with false memories (Sinclair & Barense, 2019; St. Jacques et al., 2013; Stark et al., 2010). It was indeed reported that memory misattributions (new information mistakenly attributed to an old experience) may be associated with neural reactivation of an incorrect context (Gershman et al., 2013).

Crucially, it also remains unclear how this mechanism is influenced by emotion. Negative affect can boost both recollection and familiarity (Ochsner, 2000; Rimmele et al., 2011), and more generally emotional information leads to stronger memory traces (LaBar & Cabeza, 2006; Talmi et al., 2007; Yonelinas & Ritchey, 2015) through modulations of both sensory areas (Meaux et al., 2019; Vuilleumier, 2005)and hippocampus (Dolcos et al., 2004). However, emotional memories are typically characterized by a trade-off between the item memory enhancement and context memory impairment (Adolphs et al., 2001; Chiu et al., 2013; Kensinger & Schacter, 2006; Mather & Knight, 2008). An influential account suggests that emotional information enhances recognition memory for a given item due to up-regulation of amygdala but impairs associative memory for concomitant context due to down-regulation of the hippocampus (Bisby & Burgess, 2017).

Accordingly, content-specific reactivation in visual cortex during correct retrieval is increased for items previously paired with emotional cues, whereas brain areas responsible for contextual integration such as retrosplenial and associative cortices show decreases (Hofstetter et al., 2012). Moreover, emotion also plays an important role in post-traumatic stress disorder, where involuntary retrieval of traumatic events is associated with vivid sensory details (Brewin et al., 2010). Hence, it is possible that emotional cues might promote sensory cortical reactivations, with or without conscious recall. Alternatively, emotional signals might favour false memories and induce reactivations without veridical memory traces, due to reduced encoding or impaired retrieval of contextual information in emotional conditions (Windmann & Kutas, 2001).

To resolve these issues, we designed a memory study comprising a behavioural encoding session and a retrieval session in fMRI. Most neuroimaging studies of memory errors used recognition tests for single items (Marchewka et al., 2008; Rissman et al., 2010) or cued recall with word associates (Kahn et al., 2004; Wheeler & Buckner, 2003). Activity in lateral parietal and inferior frontal cortices was found to reflect subjective “oldness” of memories rather than true encounter history (Jaeger et al., 2013; Kurkela & Dennis, 2016), and both common and distinct neural correlates were demonstrated in false memory and deception (Lee et al., 2009; Yu et al., 2019). To further investigate the nature of cortical reactivation during memory errors, here we employed an associative memory paradigm with stimulus pairs made of unique scenes and a set of to-be-remembered faces. The reactivation of face information at test was assessed by comparing brain responses to scene cues when participants correctly vs incorrectly retrieved faces, in particular in face-selective areas of fusiform cortex (Kanwisher & Yovel, 2006), compared to a control place-responsive region in parahippocampal cortex. Critically, we also tested for emotional modulation of memory reactivation during memory errors, including misses, false alarms, and misattributions.

## 2 Material and methods

### 2.1 Participants

Twenty healthy participants (3 left-handed, 13 females, 7 males, age range 18-35, M = 25.73, SD = 5.30) participated in our experiment and gave written informed consent according to the Ethics Committee regulation of Geneva University Hospitals and Medical School. Two participants were excluded due to insufficient task performance (lower than chance level), leaving a group of 18 (3 left-handed, 13 females, 5 males, age range: 18-33 (M = 24.87, SD = 4.80).

### 2.2 Stimuli

A subset of 170 scenes (85 neutral and 85 negative) was randomly selected for each participant from a bigger set of 120 neutral and 120 emotional scenes used in a previous study (Hofstetter et al., 2012). Overall, this set of scenes had been compiled from various sources, including IAPS (Lang et al., 2008) and a dataset used and validated in our previous studies (Hofstetter et al., 2012). These scenes had various contents (objects, streets, houses, landscapes, etc.) but were devoid of faces, humans, and text. They were rated for emotional parameters (valence, arousal, and dominance, each on a scale from 0 to 100; see: (Bradley & Lang, 1994), by a group of 10 independent judges (not included in the imaging session). The average values of emotional parameters for the 120 neutral scenes were as follows: valence = 59, arousal = 44; and for the 120 negative scenes: valence = 29, arousal = 65. The negative scenes were rated significantly lower on the scale of valence (i.e. more negative) [t(238) = 36.3, p < .001, Cohen’s d = 4.69] and significantly higher on the scale of arousal [t(238) = -29.2, p < .001, Cohen’s d = -3.77].

In addition, 4 colour photographs of neutral faces (young woman, young man, old woman, old man) were selected from the Lifespan Face Database (Minear & Park, 2004; https://pal.utdallas.edu/facedb/). These served as memory targets shared across trials, to be associated with unique scene cues. Pilot testing showed that using a limited number of faces allowed us to efficiently balance the correct and incorrect memory retrieval at test.

For each participant, scenes were randomly assigned to the “old” scenes presented during encoding or “new” scenes serving as lures during retrieval test. Moreover, for each participant, faces were randomly paired with different study scenes, with a constraint that each face was associated equally often to negative and neutral scenes.

### 2.3 Procedure

To increase the rate of forgetting and possibly increase emotion effects on memory due to consolidation (see: Yonelinas & Ritchey, 2015), the study took place on two consecutive days. On day one, the encoding phase was performed in front of a computer screen in a quiet room. On day two (delay in hours: M = 19.93, SD = 4.30), a retrieval task was performed while undergoing functional magnetic resonance imaging (fMRI). In addition to the memory task, participants underwent a structural anatomical scan and a functional localizer scan to delineate face responsive regions. All visual stimuli were presented on a back-projection screen inside the scanner bore using an LCD projector (CP-SX1350, Hitachi, Japan). Responses were recorded with a response button box (HH-1×4-CR, Current Designs Inc., USA).

#### 2.3.1 Memory Task

The encoding phase of the experiment comprised the presentation of 120 scene-face pairs, where one of the four faces (i.e. identities - young woman, young man, old woman, old man) was presented above a unique scene, either neutral or emotional, for 4 s (Fig. 1A). A blank screen was shown for 1 second between each trial. Each scene-face pair was presented once only, and participants were instructed to memorize the pairs; no behavioral response was requested at encoding. During the retrieval phase that took place the next day in the MRI scanner, participants were sequentially presented with the scenes and asked to indicate which face was previously associated with each scene (or whether it was new; see Fig. 1B). This phase comprised 170 scenes (120 old, 50 new; ∼11.9° x 8.4° degrees), each presented for 4.5 s, followed by a blank screen of variable duration (1.1-4s). Together with the scene, five boxes were presented on the screen as response options. The boxes had the following five labels (translated to English): “young woman”, “young man”, “old woman”, “old man”, and “new scene”. The order of the labels was changed in the middle of the experiment. At the beginning of each trial, the box in the center was marked as “selected” (highlighted with grey frame), whereas the other labels were “deselected” (blue frames). On the response box, by using their index finger and middle finger, the participants had to move the grey frame to select the appropriate response, and then confirmed their selection with pressing a third key with the ring finger (causing the chosen box frame to turn black; see Fig. 1).

**Fig 1.**
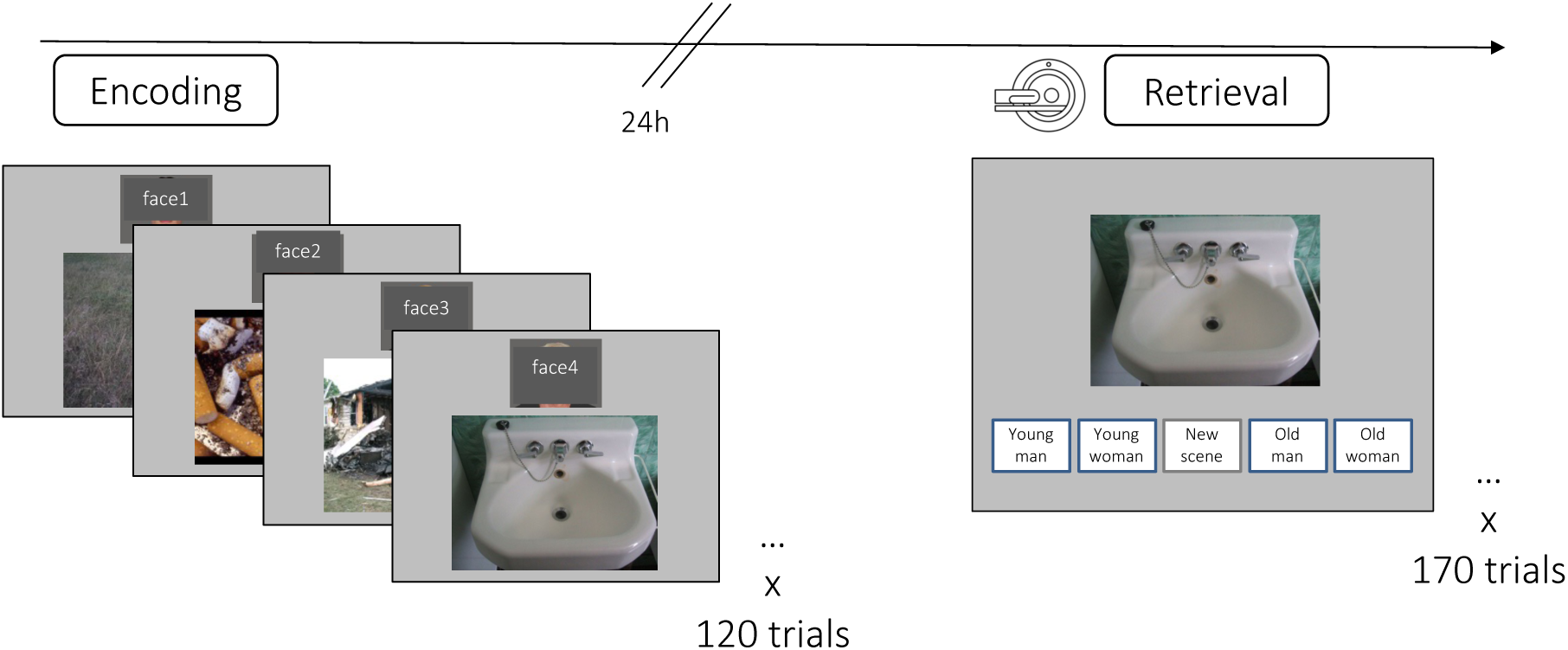
Overview of experimental procedure with two phases: Encoding - participants studied a series of 120 face-scene pairs, where each scene was associated to one of four associated faces: young woman, young man, old woman or old man (50% of the scenes was negative, 50% was neutral); Retrieval – on the following day, during an fMRI session, participants performed a scene-cued associative retrieval task, where they had to recognize whether a unique scene (n = 170) was “old” (previously studied) or “new” (not studied before) and to retrieve one of the four faces associated with the scene during encoding (50% of the cue scenes was negative, 50% was neutral).

**Fig 2.**
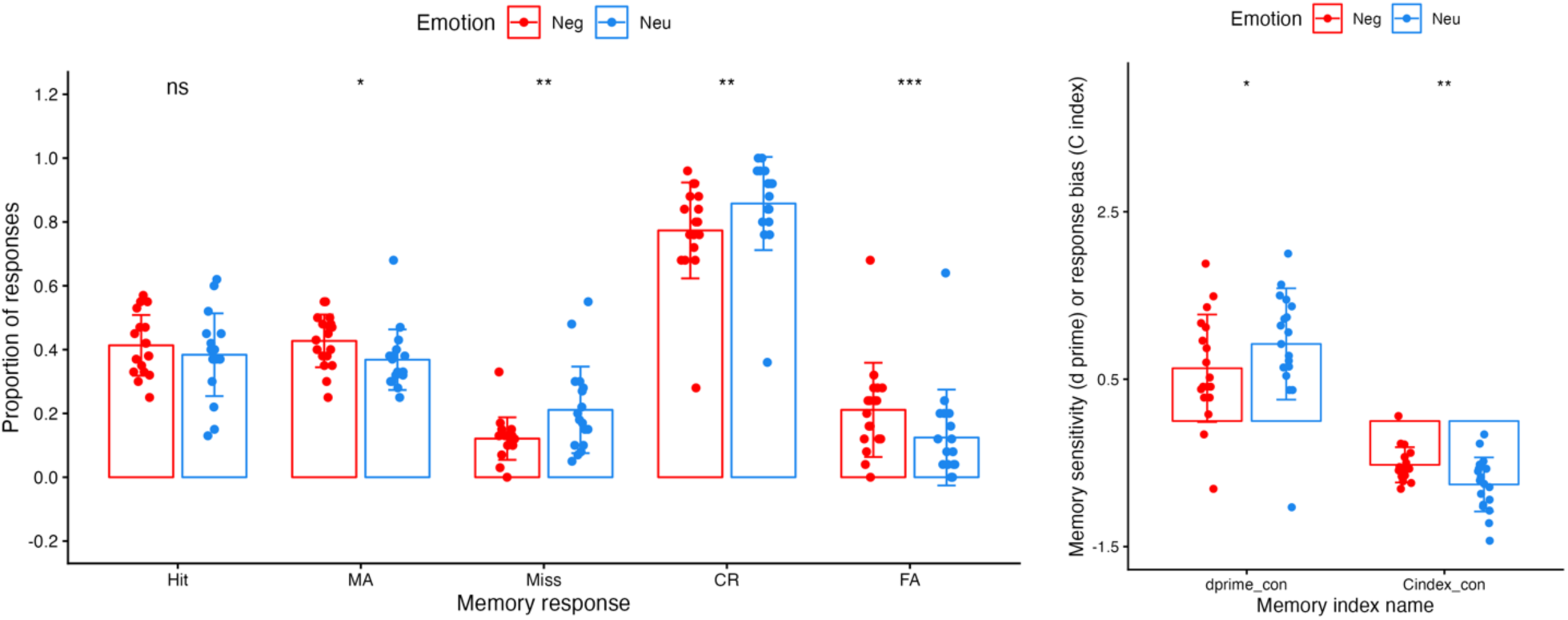
**A**. Proportion different memory responses given in the retrieval phase. Emotional content of scene cues had a significant effect on the MA, Miss, CR, and FA response rate. Note that memory performance (Hit+CR/#trials) was significantly above chance level for all participants included in our analysis (n=18). Error bars depict the standard error of the mean. **B.** Memory sensitivity index (d prime) and response bias (C index) as a function of emotion. The indices were calculated in a conservative way (considering only Hits and no MAs as correct responses, as described above). MA - Misattribution, CR - correct rejection, FA - false alarm]. *p < .05; ** p < .005; *** p < .001

#### 2.3.2 Functional Localizer Task

The localizer comprised 4 blocks of 4 different stimulus classes, with 16 images in each block (650 ms + blank 433 ms per trial). The visual stimuli were grey-scale pictures of faces, houses, words, as well as random texture patterns of V letters (Hofstetter et al., 2012). To ensure a sustained attention and similar task demands, subjects had to press a button upon the presentation of a rare upside-down item (6% of trials for each stimulus category). The mean accuracy of performance in this task (hit rate – false alarm rate) across participants was 97%. A fixation cross was shown for 4.3s between blocks.

### 2.4 Image Acquisition

MRI images were acquired on a 3T whole-body MRI scanner (Trio TIM, Siemens, Germany) with a 12-channel head coil. Functional images for BOLD contrast were acquired with a susceptibility weighted EPI sequence (TR/TE = 2100/ 30 ms, flip angle = 80 degrees, PAT factor = 2, 64 x 64-pixel, 3.2 x 3.2 mm, 36 slices, 3.2 mm slice thickness, 20% slice gap). The structural images for anatomy were acquired with a T1 weighted 3D sequence (MPRAGE, TR/TI/TE = 1900/ 900/ 2.27 ms, flip angle = 9 degrees, voxel dimensions: .9 mm isotropic, 256 x 246 x 192 voxel).

### 2.5 Data analysis

#### 2.5.1 Behaviour

First, we calculated a retrieval rate by dividing all confirmed correct responses (i.e. Hits and Correct Rejections given by the participants within the trial duration) by a total number of trials (i.e. 170). Of note, the chance level in this task was 20%, given 5 possible responses.

Then, we ran an additional analysis to control for possible differences in the level of uncertainty of memory decisions, and the resulting strategies of responding. This analysis was based on the signal detection theory (SDT; Kellen et al., 2021; Swets et al., 1978; Wickens, 2002). We calculated the proportion of: Hits, i.e., correctly retrieved face associated with an old scene (“Old” | Target); Misattributions, i.e., incorrectly retrieved face in response to a correctly recognized old scene (incorrect “Old” | Target); False Alarms, i.e. new scenes incorrectly recognized as old (“Old” | Foil), Correct Rejections, i.e., new scenes correctly retrieved as new (“New” | Foil) and Misses, i.e., old scenes incorrectly recognized as new (“New” | Target). Based on these proportions, we calculated: i) *sensitivity index (d’)* to provide a measure of the separation between the means of the signal and noise distributions, compared against the standard deviation of this distribution: Z (Hit rate + Misattribution rate) – Z (False Alarm rate); and ii) *response bias* or *decision criterion (C)* to provide a measure of the general tendency to respond yes or no, as determined by the location of the criterion, such that positive values indicate liberal response bias: (Z (Hit rate + Misattribution rate) + Z (False Alarm rate)) / 2. The *d’* and *C* indices were calculated separately for each emotion category and compared in a rm ANOVA with emotion (2 levels: neutral, emotional) as a within-subject factor.

We also systematically compared retrieval performance as a function of the emotional content (negative vs. neutral) and type of memory responses given to old cues (Hits, MAs, Misses) and new cues (CR and FA). Here, we analyzed the retrieval performance in a repeated-measures ANOVA (rm ANOVA) as a function of emotion (2 levels: neutral, emotional) and a response type as within-subject factors.

In addition, we followed recent work indicating that reaction times provide similar information to explicit ratings of memory confidence in order to quantify recognition decisions (Weidemann & Kahana, 2016) and can be more informative than memory responses to understand memory-based decisions as a single or dual process (Kraemer et al., 2021). Accordingly, we performed three rm ANOVAs for a) RT of the first button press (as a proxy of initial memory access), b) RT of the second button press (as a proxy of confirmatory memory decision), and c) the RT difference between the second and first button press (as a proxy of a mnemonic decision process). In these rm ANOVAs, we included emotion (2 levels: neutral, emotional) and memory response (5 levels: Hits, MA, Miss, CR, FA) as within-subject factors.

#### 2.5.2 Functional and anatomical region-of-interest localization

The functional localizer data was analysed at the subject level using the GLM approach with 5 regressors corresponding to: faces, words, houses, texture patterns of V letters, and button presses. Face-responsive areas were determined at the group level by identifying voxels differentially activated in the contrast “faces vs. [houses, words, patterns]”. This contrast produced robust activation in bilateral fusiform cortex, stronger in the right hemisphere (see: Supplementary Table S1). Activation maps from this contrast were masked inclusively [p = .005] by the anatomical region of fusiform gyrus specified according to the AAL2 (Rolls et al., 2015). For the resulting activation maps, a voxel-wise height threshold of p < .001 (uncorrected) was combined with a cluster level of k = 10 and used as a functionally-defined region of interest (ROI) for left and right fusiform face area (FFA), see: Supplementary Fig. S2.

As a control, place-responsive areas (Epstein & Kanwisher, 1998; Li et al., 2022) were determined at the group level by identifying voxels differentially activated in the contrast “houses vs. faces, words, patterns”. This contrast produced robust activation in bilateral parahippocampal cortex (PPA), stronger in the right hemisphere (see: Supplementary Table S1). Activation maps from this contrast were masked inclusively [p = .005] by the anatomical region of the parahippocampal cortex according to the AAL2 (Rolls et al., 2015). For the resulting activation maps, a voxel-wise height threshold of p < .001 (uncorrected) was combined with a cluster level of k = 10 and used as a functionally defined region of interest (ROI) for left and right parahippocampal place area (PPA). see: Supplementary Fig. S2.

Based on the available literature, several regions of interest (ROI) were selected to further test specific hypotheses concerning differences in emotional modulation of successful retrieval and memory errors. Anatomical masks of bilateral amygdala (AMY) and hippocampus (HC) were specified according to the AAL2 (Rolls et al., 2015) atlas implemented in the WFU PickAtlas (Maldjian et al., 2003) toolbox, version 3..5. Functional masks (FFA and PPA) were defined based on the results of a functional localizer task, described above. The ROI analysis was performed using a MarsBaR toolbox (Brett et al., 2002). Contrast estimate values were extracted and averaged based on the subject-level SPM models used in the first univariate ANOVA analysis mentioned below in the following regions.

#### 2.5.3 Region-of-Interest analyses

To investigate the effects of emotion and memory response in these ROIs during the retrieval phase, we performed rm ANOVAs on the contrast estimates from each of the ROIs: FFA left and right (reported in the manuscript), PPA, AMY, HC left and right (control regions, reported in Supplementary materials, Fig. S3). We analyzed the contrast estimates as a function of two within-subject factors: memory response (5 levels: Hits, MA, Miss, CR, FA) and emotion (2 levels: negative, neutral). Second, we performed a post hoc analysis of simple effects, namely a paired t-test for dependent samples to directly compare the mean contrast estimates corresponding to different conditions, based on a priori hypotheses.

One subject did not have any Miss responses for negative scene cues and three participants did not have FA for negative and neutral scene cues; hence for the corresponding ANOVA we substituted the missing values with the condition group mean (Vaden et al., 2012). Excluding these participants from the analyses did not change the outcome of the ANOVA, see: Supplementary materials and (Pine et al., 2018) for a similar approach.

#### 2.5.4 Univariate whole-brain activation analysis

All pre-processing and statistical analyses of imaging data were performed with the Statistical Parametric Mapping (SPM12; Wellcome Department of Cognitive Neuroscience, University College London, London, UK) running under Matlab 2013b (Mathworks, Inc., Natnick, MA, USA), using standard procedures. The pre-processing included realignment, slice timing correction, normalisation, and smoothing (8 mm FWHM Gaussian kernel).

Then, first-level analyses were conducted for each subject within the general linear model (GLM) approach. Each single event was modelled with onset and duration corresponding to the presentation of the scene cue, until the response button press. The model included 12 regressors for experimental conditions: Hit (retrieval of correct face), Misattribution (retrieval of another, incorrect face), Miss (responding new to scenes previously associated with a face), Correct Rejection (correctly responding new), False Alarm (incorrectly recalling a face in response to a new scene), and rare (3%) Unconfirmed trials (no final key press during trial duration) - each of these memory responses for Negative and Neutral cues. To account for movement-related variance, six nuisance regressors were included as representing the differential of the movement parameters from the realignment. Data was high-pass filtered (1/128 Hz) and convolved with a standard canonical hemodynamic response function (HRF) to approximate the expected blood-oxygen-level dependent (BOLD) signal.

The second-level (random-effect) whole brain group analysis was performed using ANOVAs evaluated to obtain estimates of activity in response to each trial type relative to implicit baseline. Flexible factorial design (Ashburner et al., 2014) was used because of the abovementioned variability in the number of trials among subjects and possible subject effects. In this analysis, the interaction between emotion and memory performance was examined for the retrieval data. Functional volumes from encoding session were split into conditions along the factors of emotion: Negative (Neg), Neutral (Neu) and subsequent memory performance: Hit, MA (Misattribution), Miss, CR (Correct Rejection), FA (False Alarm). Thus, 10 experimental conditions were specified as follows: Neg FA, Neg MA, Neg Miss, Neg CR, Neg FA, and Neu FA, Neu MA, Neu Miss, Neu CR, Neu FA. The number of trials in each condition varied according to subsequent individual memory performance, but all subjects showed overall hit rates above chance level (see results).

Whole-brain random-effects contrasts were evaluated to obtain estimates of activity in response to each trial type relative to implicit baseline. Event-related stick-function regressors were used to perform ANOVA with emotion (two levels: Neg, Neu) and memory (5 levels: Hit, MA, Miss, CR, FA) as a within-subject factors, as well as a subject factor.

Unless stated otherwise, we applied a voxel-wise height threshold of p < .001 (uncorrected) combined with a cluster size of k = 10 in the whole brain analyses. The small volume correction (svc.) (Worsley et al., 1996) was applied using anatomical masks corresponding to the regions-of-interest (ROI) selected based on a priori hypotheses about their engagement in emotion, memory and sensory cortical regions interactions: FFA (reported in the manuscript), PPA, AMY, HC (reported in supplementary materials, Fig. S3). Within each of these ROIs, we considered activations whose effects survived the small volume FWE correction at the voxel level. This type of correction for multiple comparisons was previously applied in studies of emotion-memory interactions due to a small volume of interest (Dominiguez-Borras et al., 2017; Richter et al., 2016).

The coordinates of significant effects were reported in the Montreal Neurological Institute (MNI) space and labeled according to Automated Anatomical Labeling (AAL2) (Rolls et al., 2015) atlas with the use of bspmview (https://www.bobspunt.com/software/bspmview/). We used MRIcroGL (http://www.mccauslandcenter.sc.edu/mricrogl/home), SPM and xjView (http://www.alivelearn.net/xjview8/) to visualize our results.

### 3 Results

### 3.1 Behavioral results

Memory of faces associated to scene cues was tested one day after encoding. All the participants included in the analyses (n = 18) performed the retrieval task above chance level (i.e. 20% - given 5 possible responses). The average retrieval rate, calculated as ([Hit + CR]/ number of trials), was 52% (SD = .08) and significantly different from zero [t(17) = 26.6; p < .001].

To consider possible responding strategies of the participants, we also computed a sensitivity index (d’) and a response bias (C index) derived from signal detection theory (SDT; Kellen et al., 2021; Swets et al., 1978; Wickens, 2002). For each participant, a normalized mean error rate (FAs) was subtracted from a normalized correct retrieval rate (Hits + MAs) to calculate d’. This revealed that the mean memory sensitivity index d’ (M = 1.93, SD = 46) was significantly different from zero [t(17) = 17.9; p < .001]. Of note, when calculated in a conservative way (only Hits – FAs, i.e., without trials where an incorrect face was retrieved in response to an old scene), the mean memory sensitivity index d’ (M = .75, SD = .60) was still significantly different from zero [t(17) = 5.30; p < .001]. We also computed the C index from the normalized mean error rate (FAs) and correct retrieval rate (Hits + MAs) of each participant. The mean response bias C index (M = -.03, SD = .33) was not different from zero (t(17) = -.38; p = .71). However, a conservative calculation of the C index (using only Hits – FAs) revealed the mean response bias (M = -.62, SD = .23) was significantly different from zero (t(17) = - 11.56; p < .001). This mild bias observed when excluding MAs accords reflects a tendency to rely on familiarity-driven responses to old scenes, with greater difficulty to retrieve the correct face associate. Overall, these results show that participants were able to remember the correct face-scene associations in a reliable proportion of trials, even though the task was purposefully designed to produce a large number of errors.

Next, we aimed at comparing these measures of memory performance as a function of emotion. To this end, we calculated d’ separately for negative scene cues (liberal: M = 1.99, SD = .65; conservative: M = .63, SD = .64) and neutral cues (liberal: M = 2.02, SD = .44, conservative: M = .92, SD = .76), and C index separately for negative scene cues (liberal: M = .16, SD = .44, conservative: M = -.52, SD = .21) and neutral cues (liberal: M = -.21, SD = .44, conservative: M = -.76, SD = .32). We found that the conservative measure of memory sensitivity (d’ calculated based on Hits - FAs) was significantly modulated by emotion [t(17) = -2.6, p = .019], but this was not the case for a more liberal measure (d’ calculated based on Hits + MAs - FAs) [t(17) = -.22, p = .83]. Specifically, memory sensitivity was higher when cued with neutral compared to negative scenes (p = .019). For response bias, we found that the liberal index (C, calculated based on Hits + MAs - FAs) differed across emotion conditions [t(17) = 5.48, p < .001], being lower for neutral than negative scene cues (p < .001). This was also the case for the conservative index (C, calculated based on Hits - FAs) [t(17) = 3.77, p = .002]. This result indicates a greater bias to respond “old” when cued with negative scenes.

We also systematically compared retrieval performance as a function of the emotional content (negative vs. neutral) and type of memory responses given to old cues (Hits, MAs, Misses) and new cues (CR and FA). For old scene cues, a repeated-measure ANOVA over these two factors (emotion and response type) showed no effect of emotion [F(1,17) = .01, p = .897, η^2^ = .000], but a significant effect of response [F(2,34) = 31.71, p < .001, η^2^ = .517] and, critically, a response type x emotion interaction [F(2,34) = 4.82, p = .019, η^2^ = .045]. Follow-up paired t-tests showed that participants significantly more often gave MA responses to negative than neutral cues [t(17) = 2.38, p = .03] (although this result did not survive a correction for multiple comparisons, p = .44, Bonf. corr.). On the contrary, participants gave significantly less Miss responses to negative cues than to neutral cues [t(17) = -3.35, p = .004; p = .058, Bonf. corr.]. There was no significant difference in Hit responses to negative and neutral cues [t(17) = .86, p = .40; p = 1.00, Bonf. corr.]. For the new scene cues, a paired t-test showed significantly more CR responses to novel neutral (M = .86, SD = .15) than to novel negative cues (M = .77, SD = .15) [t(17) = -4.86; p < .001]. Conversely, as implied by this result (since CR = 1 - FA), FA responses were more frequent to novel negative cues (M = .21, SD = .15) than to novel neutral cues (M = .712, SD = .15) [t(17) = 5.96; p < .001].

To better understand temporal dynamics of making a memory decision after an initial memory access, we also analyzed the reaction time (RT) on the scene-cued associative retrieval task as a function of memory response and emotion. Specifically, we examined a) RTs for the first button press - response choice, as a proxy of initial memory access, b) RTs for the second button press – response confirmation, as a proxy of memory decision, and c) RT differences between the two button presses as a more precise timing of the memory decision after initial memory access.

We found that RT for the first button press was modulated both by emotion and memory response (see: Supplementary Fig. S1). Participants spent longer on selecting the response when cued by emotional compared to neutral scenes [F(1,10) = 12.033, p = .006, η^2^ = .546], and longer on memory errors than correct retrieval [main effect of memory response F(4,40) = 6.625, p = .003, η^2^ = .399]. Specifically, CRs were faster than FAs [t(10) = -3.54; p = .005], MAs [t(10) = -2.883; p = .016], and Misses [t(10) = -2.429; p = .035]; whereas Hits were faster than FAs [t(10) = 3.706; p = .004] and marginally faster than MAs [t(10) = -2.126; p = .059]. FAs were also slower than Misses [t(10) = 2.935; p = .015]. There was no interaction between these two factors [F(4,40) = 1.707, p = .146, η^2^ = .663].

On the other hand, RT for the confirm press was modulated by emotion [F(1,13) = 5.35, p =.038, η^2^ = .292], memory response [F(4,52) = 15.417, p < .001, η^2^ = .543], and importantly, an interaction between emotion and memory response [F(4,52) = 3.243, p = .04, η^2^ = .2]. Specifically, it took longer to confirm their response when participants were cued by emotional compared neutral scenes on trials with CRs [t(13) = 2.768, p = .064] and MAs [t(13) = 3.231, p = .033].

Finally, the duration of memory decision (RT difference between the first and confirmatory press) was modulated only by memory response [F(4,52) = 7.262, p = .003, η^2^ = .358]. There was no emotion [F(1,13) = .021, p = .866, η^2^ = .002] or interaction effect [F(4,52) = .606, p = .566, η^2^ = .045]. Participants were the fastest to confirm their initial memory response on CR – faster than Hit [t(13) = -5.023, p = .002] and MA [t(13) = -4.498, p = .005]. However, decision RTs for Hits [t(13) = 3.404, p = .005] and MAs [t(13) = 3.338, p = .005] were slower than Misses.

In sum, these behavioural results show that our paradigm successfully produced a balanced distribution of correct and incorrect recall, without any major response biases. Notably, emotional scenes were associated with fewer response omissions (Misses), but also with a greater tendency to make FA errors (incorrectly respond “old” to the new scenes) and MA errors (recalling an incorrect face paired with an old scene).

### 3.2 Imaging results

We analyzed successively memory-related activations, followed by a comparison between explicit and implicit effects, and finally emotional influence of negative vs neutral cues. For all three steps, we report specific ROI analysis for anatomically (AMY and HC) and functionally (FFA and PPA) defined regions based on a priori hypotheses, as well as a broader whole brain analyses.

#### 3.2.1 Global memory-related activations

##### 3.2.1.1 Whole brain analysis

To identify brain regions recruited during correct memory retrieval of previously seen face-scene pairs, we compared activity during Hit vs CR trials (pooling emotional and neutral conditions together). This contrast showed a widespread network consisting of precuneus, bilateral inferior parietal cortex, left inferior frontal gyrus and insula, anterior cingulate cortex, bilateral caudate, as well as left fusiform cortex (Fig. 3A and Supplementary Table S2). Activity in left fusiform overlapped anatomically with face-selective regions identified in the face localizer (svc. using the FFA activation volume functionally defined by the localizer showed that this activation survived a correction for multiple comparison; see Fig. 3B and below). This fronto-parietal network is consistent with typical findings during memory retrieval and memory vividness in other studies(Eldridge et al., 2000; Kahn et al., 2004; Wheeler & Buckner, 2003), accompanied by a recruitment of visual cortical areas engaged by the remembered stimulus category (Hofstetter et al., 2012).

**Fig. 3.**
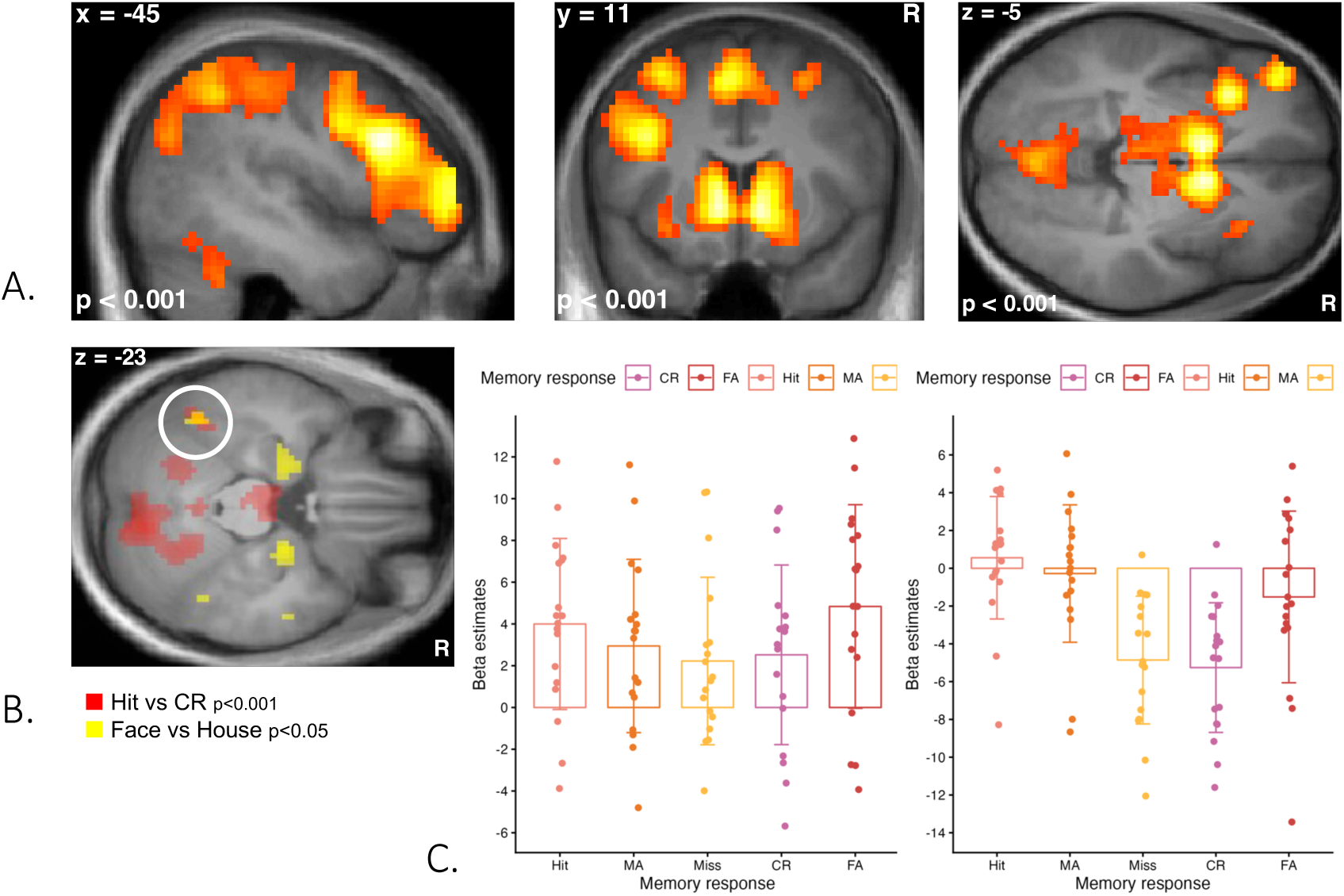
Comparison of Hits and CRs. **A.** An extensive network of brain regions showed stronger activation for Hits vs CRs including left prefrontal and parietal cortex, caudate, and visual areas overlapping with functionally defined FFA, as shown in **B** (white circle). **C.** Beta estimates across the different memory responses for peak activations in left fusiform (left panel) and precuneus (right panel), pooling over both emotion conditions. MA - Misattribution, CR - correct rejection, FA - false alarm].

Given our main focus on visual reactivation across memory conditions, we extracted beta parameters from the peak of the left fusiform cluster identified above to examine the pattern of relative activations across conditions. As illustrated in Fig. 3C, this region did not only activate during Hits (i.e. successful retrieval of the face previously associated with the scene), but also during False Alarms (in which participants saw a new scene but incorrectly reported remembering an associated face). By contrast, activity was reduced during Misses (i.e. old scenes which participants did not recall), to the same level as that observed during CR (i.e. new scenes correctly recognized as such). A similar activation pattern across conditions was observed for the precuneus (Fig. 3C), a region commonly engaged during memory retrieval (Cavanna & Trimble, 2006; Flanagin et al., 2023; Hebscher et al., 2020; Ritchey & Cooper, 2020). These data suggest that fusiform activity followed *subjective* memory responses of the participant, rather than the *objective* history of past associations between faces and scenes. Misattribution errors (MA), in which participant recognized the scene but reported an incorrect face associate, evoked somewhat intermediate activation levels in left fusiform, slightly reduced in comparison to successful retrieval (Hits), but higher than Misses. These findings were further corroborated by statistical analysis conducted on a priori defined ROIs in fusiform cortex (see next section).

##### 3.2.1.2 Fusiform ROI analysis

To specifically examine memory-related activation in fusiform cortex, we performed a region of interest (ROI) analysis restricted to voxels with face-selective activity (FFA) as defined at the group level from a separate functional localizer scan (see: Supplementary Table S1). Beta estimates extracted from FFA in both hemispheres were analyzed by ANOVA with the factors of emotion, memory response, and the interaction of emotion and memory response (reported below in section 3.2.3.2). This analysis showed a main effect of memory response [F(4,68) = 3.89, p = .043, η^2^ = .029], but no effect of emotion [F(1,17) = 2.157, p = .16, η^2^ = .004], and no interaction [F(4,68) = 2.075, p = .121, η^2^ = .009]. Consistent with our whole-brain results above, post-hoc comparisons for the left FFA ROI showed significantly higher activation for Hits compared to CR [t(17) = 5.057, p = .001], Miss [t(17) = 4.521, p = .003], and MA [t(17) = 3.5, p = .022], However, Hit did not differ from FA [t(17) = - .427, p = .675]. Activations during FA were marginally stronger than during CR and Miss, but these effects did not survive a correction for multiple comparisons [t(17) = 2.062, p = .384 and t(17) = 2.02, p = .384, respectively]. MA activation overall was of intermediate magnitude, between Hit and Miss/CR (see section 3.2.3.2 for further analyses memory response as a function of emotionality of cues).

The same analysis for right FFA showed no significant effect of memory response [F(4,68) = .748, p = .479, η^2^ = .005], no effect of emotion [F(1,17) = 1.433, p = .248, η^2^ = .005], and no interaction [F(4,68) = 2.066, p = .143, η^2^ = .029].

#### 3.2.2 Comparison of explicit and implicit memory

To probe for any implicit memory effects during retrieval, we performed a whole-brain contrast comparing Misses vs Correct Rejections (where scenes were new and never paired with faces). This contrast revealed clusters in bilateral posterior parahippocampal cortex (PHC; xyz = -18 -52 -5, Z = 3.93; xyz = 21 -49 -8, Z = 4.14), insula (xyz = -30 17 -11, Z = 3.71), and anterior cingulate cortex (xyz = - 3 17 49, Z = 3.43; all p < .001), consistent with some effect of covert recognition or uncertainty during Misses (Fig. 4A). However, there was no effect in fusiform cortex or any other visual areas, even at lower thresholds.

**Fig. 4.**
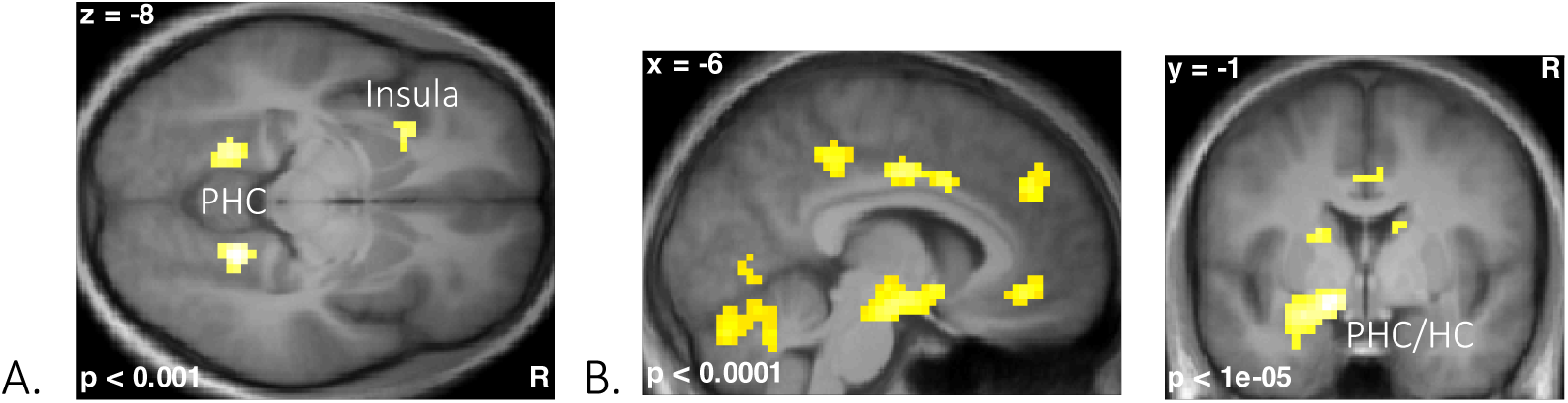
Implicit and explicit effects. **A.** Implicit effect: The comparison of Miss vs. CR trials produced activations in bilateral parahippocampal gyri and left insula. **B.** Explicit effect: Comparing Hit vs MA yielded a network of activations including ventromedial prefrontal cortex (VMPFC), posterior cingulate cortex (PCC), anterior cingulate cortex (ACC) and left hippocampus (HC) and parahippocampal cortex (HPC).

Conversely, we determined brain regions underlying explicit retrieval by contrasting Hits to MA. In both conditions, participants reported remembering a face associated with the scene, which was either the correct face (Hit) or an incorrect face (MA). This analysis indicated that Hits engaged a distinctive network (Fig. 4B), involving the hippocampus and parahippocampal cortex (PHC; xyz = -24 -25 -17, Z = 4.27), as well as the ventromedial prefrontal cortex (VMPFC; xyz = -9 35 -11, Z = 5.60), the rostral (xyz = 0 47 -2, Z = 4.42) and posterior cingulate (xyz = -3 -34 40, Z = 4.17), the inferior parietal lobule (xyz = -51 -58 16, Z = 3.92), plus lateral temporal cortices (xyz = -57 -34 -8, Z = 4.55; xyz = 63 - 34 -8, Z = 4.25; all p < .001). This hippocampal-prefrontal cortical network has often been implicated in episodic or autobiographic memory (Cabeza et al., 2004; Fink et al., 1996). In addition, this contrast also revealed significant activation in extrastriate visual areas that overlapped with face-responsive ROIs from the localizer scan (left FFA peak, xyz = 42 -49 -23, Z = 2.8, p = .003). Taken together, these data converge to indicate that face-specific reactivation in visual cortex occurred under explicit memory conditions, without any evidence for implicit effects in this region.

Finally, when examining the functionally defined FFA ROIs, we did not find any covert modulation in response to scenes previously paired with a face when participants did not retrieve this association.

#### 3.2.3 Emotional modulations

##### 3.2.3.1 Whole brain analysis

The main effect of emotion was first tested by comparing negative vs neutral scene cues across all memory conditions. This revealed an expected pattern of activations in visual areas (xyz = 30 -76 - 11, Z = 6.12; xyz = -24 -88 -14, Z = 5.87), amygdala (xyz = -21 -4 -23, Z = 5.04; xyz = 24 -1 -26, Z = 3.50), orbitofrontal cortex (xyz = -33 32 -17, Z = 4.83; xyz = 33 32 -20, Z = 4.30), and parahippocampal cortex (xyz = -21 -25 -11, Z = 3.92; all p < .001), confirming the significant emotional impact of these pictures.

When considering Hits alone, we found no differential increase in fusiform regions when the to-be-remembered faces had been paired with a negative (vs neutral) scene, suggesting that emotional cues did not reliably modulate cortical reactivation in visual areas during successful retrieval. This was further confirmed by the contrast Hit > CR for each emotion condition, which revealed highly overlapping activations in fronto-parietal networks, precuneus, insula, and caudate (see Fig. 6).

##### 3.2.3.2 FFA ROIs analysis

Given our focus on visual reactivation and a priori hypotheses about emotional modulation of brain activations across memory responses, we also inspected the functionally defined FFA ROIs from the localizer scan, now splitting each memory condition into negative and neutral trials.

As shown in Fig. 5, the left FFA showed no significant emotional effect during Hits [t(17) = -.516, p = .613], or FA [t(17) = - .44, p = .666], whereas a remarkable increase was observed for CR [t(17) = - 5.351, p < .001] and MA [t(17) = - 2.868, p = .043]. At the same time, direct comparisons with pairwise t-tests confirmed that MA in response to a negative scene cue produced similar FFA activations to Hits [t(17) = .945, p = .358], but differed from the Miss trials [t(17) = 2.44, p = .413] (yet did not survive a correction for multiple comparisons). On the other hand, MA in neutral cues evoked low activity in FFA, which differed from neutral Hits (yet did not survive a correction for multiple comparisons) [t(17) = 2.964, p = .157], and was equivalent to the Miss [t(17) = -.449, p = .659] or CR [t(17) = .828, p = .419] conditions (Fig. 5A). This pattern explains why overall MA responses showed a response intermediate between Hits and Miss or CR in our previous analysis, when pooling both emotion conditions together (see above section 3.2.1.2).

**Fig 5.**
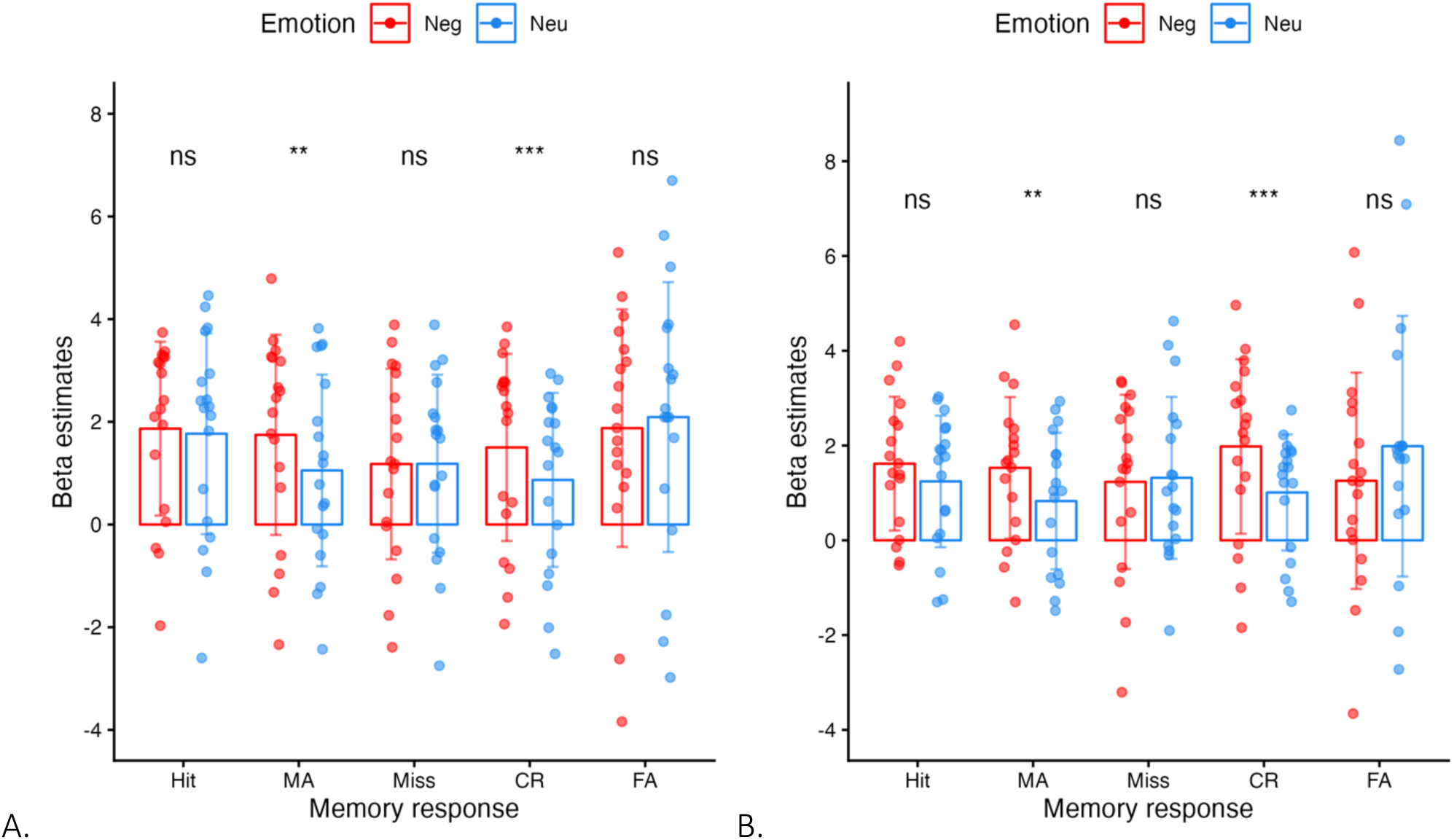
FFA was defined in both hemispheres by an independent face localizer scan. **A.** Modulation of left FFA by memory condition and emotional context. Beta values extracted from the left FFA showed general increases for Hit and False Alarms, whereas the emotional scene content produced significant modulations specifically for MA and CR. Activation of the left FFA during MA for negative emotional scenes did not differ from Hit (negative and neutral); however, MA for neutral scenes showed a much lower activation comparable to Miss (negative and neutral). **B.** Modulation of right FFA by memory condition and emotional context. We did not observe any emotional effect during Hits, but we did observe that activations to emotionally cued MA and CR were also higher in right FFA. *p < .05; ** p < .005; *** p < .001

**Fig. 6.**
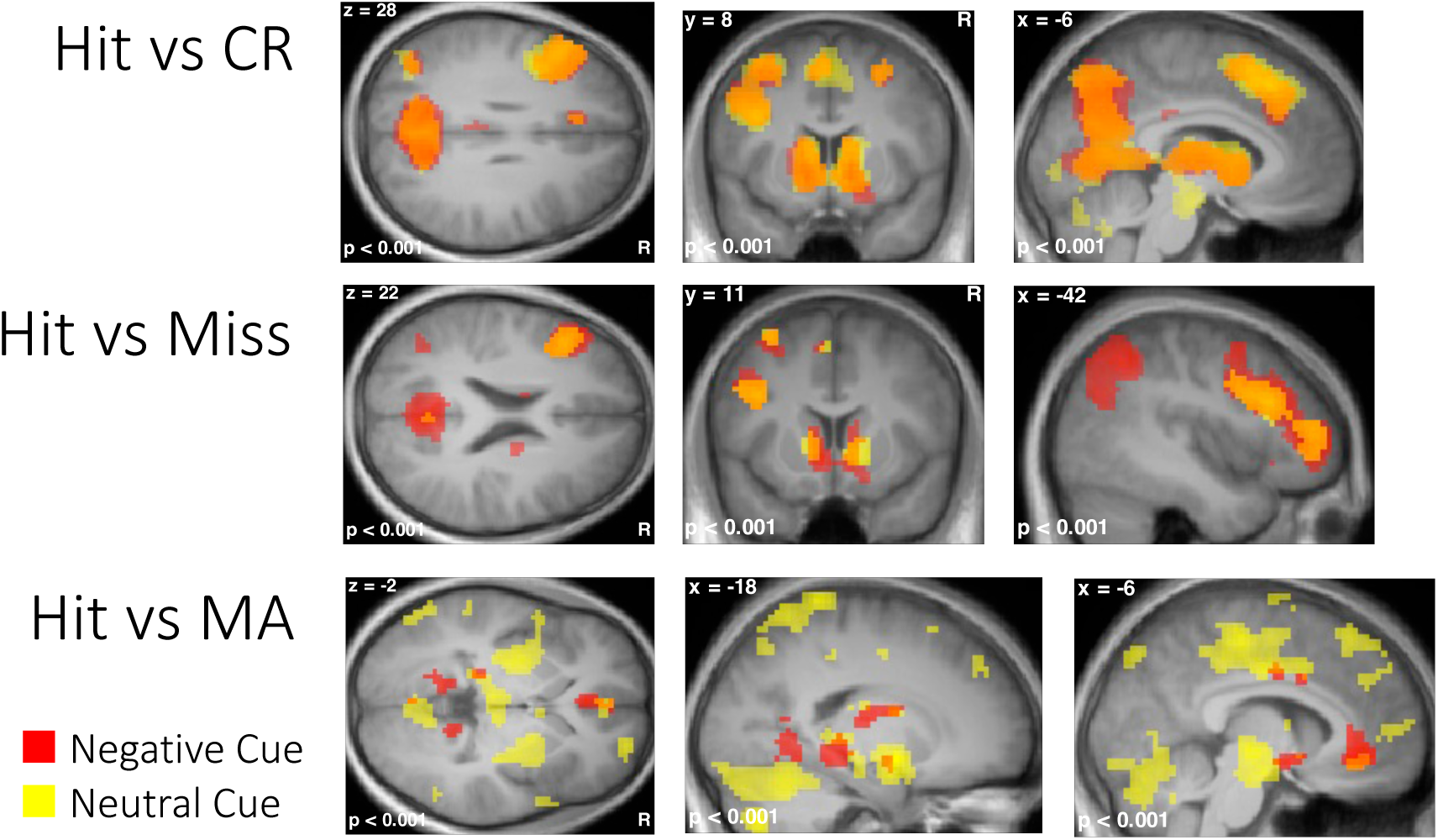
Comparison between negative and neutral cues for different memory conditions. In whole-brain contrasts, Hit vs Miss and Hit vs CR showed a similar network encompassing precuneus/retrosplenial cortex, left IPS, left DLPFC, left insula, ACC/ SMA, and bilateral caudate, with consistent overlap between negative and neutral conditions. Only MA trials showed distinctive emotional effects. Contrasting Hit vs MA in cued by negative scenes showed only a few differences restricted to VMPFC, anterior and posterior cingulate, hippocampus and parahippocampal cortex, while a much more distributed network in parietal and prefrontal areas was recruited when cued by neutral scenes.

We ran similar post-hoc analysis in the right FFA, driven by a priori hypotheses about emotional modulation across memory responses. As shown in Fig. 5B, we did not observe any emotional effect during Hits [t(17) = -1.924, p = .214], FA [t(17) = .865, p = .798] or Miss [t(17) = .205, p = .84], but we observed a significant emotion effect for CR [t(17) = -3.502, p = .014], and a trend effect for MA [t(17) = -2.528, p = .087]. No significant effects were observed when comparing across memory responses to negative scene cues, or across memory responses to neutral scene cues.

##### 3.2.3.3 Emotional enhancement of misattribution errors

To further decipher the emotion effects seen during MA, we compared these trials with other memory conditions in whole-brain analyses. When contrasting Hit vs MA (i.e., correct vs incorrect face retrieval) cued by negative scenes, we observed a highly a circumscribed network comprising the VMPFC, anterior and posterior cingulate, thalamus, amygdala, hippocampus, and parahippocampal cortex, a pattern reminiscent of the network that was differentially recruited by explicit retrieval regardless of emotion conditions (see above, Fig. 5B, Fig. 6 and Supplementary Table S3). The same comparison in the case of neutral cues showed a much more widespread network including the cingulate gyrus, midbrain, bilateral middle temporal, superior medial frontal, bilateral parietal lobe, as well as amygdala, hippocampus, and parahippocampal cortex (Fig. 6 and Supplementary Table S3).

Conversely, when comparing MA vs Miss (i.e., incorrect face retrieval vs forgetting), negative cues produced stronger and widespread effects in fronto-parietal networks associated with memory retrieval (Fig. 6), including left IPS (xyz = -33 -61 43, Z = 4.79), precuneus (xyz = 3 -64 28, Z = 5.96), left IFG (xyz = -48 23 28, Z = 6.08; xyz = -39 44 4, Z = 5.77), left insula (xyz = -33 20 1, Z = 4.34), and caudate (xyz = -9 8 1, Z = 4.93; xyz = 9 5 -5, Z = 3.46; all above p < .001); whereas the same comparison for neutral cue showed only limited increases in precuneus (xyz = 3 -64 28, Z = 3.50) and left DLPFC (xyz = -42 20 25, Z = 4.56, all p < .001).

Therefore, both whole brain and FFA ROI analyses converge to suggest that MA seen with <u>negative</u> cues implicate a similar network of activations as the Hit condition (negative and neutral), whereas MA in response to neutral cues is more similar to the Miss condition (negative and neutral). Crucially, however, regions in VMPFC and posterior parahippocampal cortex (found above when contrasting Hit vs MA) seem to be specific to true memory (i.e. Hit), and not modulated by emotional cue during MA.

#### 3.2.4 Psychophysiological interactions

Finally, we asked whether the modulations of FFA activity during both correct retrieval of faces (Hits) and erroneous retrieval of face associates (MAs) in the presence of emotional vs. neutral cues were driven by distinctive functional interactions with other brain regions implicated in memory and/or emotion processing, respectively. To this aim, we performed a psychophysiological interaction (PPI) analysis of the left FFA time-course, probing for differential increases in functional connectivity as a function of the emotional content of scenes (negative vs neutral).

When conducted on data from Hits, the PPI analysis of FFA connectivity revealed significant effects in left hippocampus (xyz = -27 -16 -23, Z = 3.70), as well as in the left putamen (xyz = -24 -16 1, Z = 3.54) and right inferior temporal gyrus (xyz = 48 -52 -14, Z = 3.66, all p < .001 see Fig. 7). This effect indicates that functional connectivity between left FFA and left hippocampus was enhanced on correct retrieval when the scene cue was emotional (see Fig. 7). The PPI analysis performed on MA did not produce any such activation at conventional thresholds (p < .001), confirming that these trials did not implicate the same memory circuits as Hits.

**Fig. 7.**
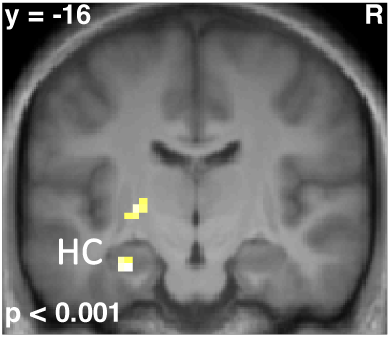
Psychophysiological interaction between of the time-course of activity in left FFA and cue emotionality during Hit. Functional connectivity of left FFA was selectively increased with left hippocampus cued by negative relative to neutral scenes.

## 4 Discussion

Memory retrieval is associated with a content-specific reactivation of sensory cortices, and such reactivation is stronger when retrieval is cued with emotional compared to neutral information. However, does similar reactivation occur during memory errors, and is it enhanced in emotional contexts? Here, we investigated the functional significance of cortical reactivation during memory retrieval and the impact of emotional cues on this process. Participants took part in a behavioural encoding session and a scene-cued associative retrieval session in fMRI on the following day. Our paradigm was specifically designed to compare behavioral performance and neural responses when participants successfully recalled a face paired with a given scene (Hits), relative to when they forgot the association (Misses) or instead reported false memories (Misattributions to another face on trials with old scenes, False Alarms on trials with new scenes). We compared the reactivation of face-selective areas of fusiform cortex, both for correct face retrieval and for memory errors. Importantly, we also tested for emotional modulation of memory reactivation during memory errors. Behaviourally, we found that emotional compared to neutral scenes induced higher rates of FA and MA, but lower rates of CR and Misses. Neurally, we found more activation in the face-specific region in fusiform cortex for both Hits and False Alarms, as compared to Misses and Correct Rejections. Critically, during Misattribution errors, we observed increased fusiform activation comparable to Hits when cued by negative scenes, but low fusiform activation similar to Misses when cued by neutral scenes. By contrast, Hits cued by emotional information recruited a distinctive network previously linked to personal episodic memories, paralleled with enhanced functional coupling between the hippocampus and fusiform cortex.

### 4.1 Validation and limitation of experimental manipulations

We validated our experimental manipulation in two ways. First, we analyzed the affective ratings provided by a group of independent judges for all the stimuli (emotional or neutral scenes) used in our experiment. As intended, negative scenes were rated as more negative and more arousing than neutral stimuli. Importantly, existing models and empirical studies point to a distinct role of negative (Bowen et al., 2019) vs. positive valence (Madan et al., 2019) on associative memory, as well as on neural reinstatement (Clewett & Murty, 2019). Here we focused on negative valence, but future studies are needed to extend our investigation to the effects of positive emotion on cortical reactivation during correct and incorrect memory retrieval.

Second, our memory task was validated by careful analysis of behavioural performance in the retrieval phase. We found that all the participants were able to reliably retrieve correct face-scene associations, even though the task was designed to produce a large number of errors. Thanks to this well-balanced task performance, we could directly compare the reactivation of face information in face-selective areas of fusiform cortex for correct retrieval and memory errors. Future studies may usefully complement this research by collecting fMRI data during both retrieval and encoding sessions. This would allow for a refine, trial-level analysis of cortical reactivation in terms of encoding-retrieval similarity for correct and erroneous memory responses, and the role of emotional context (Gershman et al., 2013; Ritchey et al., 2013).

### 4.2 Emotional modulation of memory errors

Crucially, we found that emotional scenes induced higher rates of False Alarms and Misattributions, but lower rates of Correct Rejections and Misses, relative to neutral cues. These results suggest that emotional cues induced a bias to report remembering a face, when no face or another face was seen in the past. In contrast, successful recollection of faces (Hits) was not affected by emotional cues.

This bias towards faulty memories for negative scenes might reflect subjective familiarity signals elicited by emotional content and mistakenly assigned to one of the target faces. Other studies reported that emotion can enhance the feeling of remembering even when memory is less accurate (Antypa et al., 2019; Rimmele, Davachi, Petrov, et al., 2011; Sharot & Phelps, 2004), and thus promote false alarms (Hofstetter et al., 2012; Pierce & Kensinger, 2011; Windmann & Kutas, 2001). Increased reliance on familiarity might be favored by reduced recollection of contextual information in emotional conditions; indeed, emotion often enhances memory for gist at the cost of details (Adolphs et al., 2001, 2005; Kensinger et al., 2007) or reduces the retention of precise contextual associations (Hofstetter et al., 2012; Pierce & Kensinger, 2011).

### 4.3 Content-specific, explicit vs. implicit memory reactivation

Our fMRI data revealed distinctive activation patterns dovetailing with memory performance. Successful retrieval recruited a distributed fronto-parietal network, consistent with studies on memory recollection (Eldridge et al., 2000; Kahn et al., 2004; Wheeler & Buckner, 2003), as well as face-responsive regions in fusiform cortex. The latter accords with empirical studies showing stimulus-specific reactivation during cued retrieval (Hofstetter et al., 2012; Skinner et al., 2010; Wheeler et al., 2000) or spontaneous retrieval (Polyn, 2005).

Unlike previous work, we were able to test for the role of explicit/conscious retrieval in such reactivation. To do so, we directly compared brain activity during correct retrieval with memory errors, either when recollection fails (Misses) or when a face is erroneously recalled (Misattributions or False Alarms). In the functionally-defined left FFA, we found that Hits produced the strongest activation and differed from all other conditions, except for False Alarms. In contrast, the lowest FFA activation was observed for Correct Rejections and Misses, when no face was recalled. Interestingly, Misattribution errors produced intermediate responses (when collapsing over responses to neutral and emotional cues). This overall pattern suggests that the (re)activation of cortical visual areas is primarily linked to subjective response, rather than to actual associations formed during encoding. Our results converge with recent reports (Kahn et al., 2004; Rissman et al., 2010) that activation patterns at the whole-brain level could successfully discriminate between successful recollection vs. forgetting of single items, whereas the distinction between subjective veridical and false recognition was not significant.

In addition, we found no evidence for implicit memory activation in the FFA (i.e. during the Misses), a result further supporting the link between sensory reactivation and subjective, rather than objective, memory. Implicit memory effects were observed in posterior parahippocampal cortex, possibly resulting from covert memory traces (Daselaar et al., 2006; Grunwald et al., 2003), plus in cingulate and insula, likely reflecting uncertainty or monitoring processes (van der Meulen et al., 2012).

By contrast, correct explicit retrieval (Hits vs. Misattributions) elicited a distinctive pattern in midline structures, including ventromedial prefrontal cortex, anterior and posterior cingulate cortex, as well as hippocampus and anterior parahippocampal cortex, reminiscent of results from studies on autobiographical memory (Cabeza et al., 2004; Fink et al., 1996). Likewise, brain activations in the hippocampus and parahippocampal cortex were previously observed during veridical recollection when comparing “remember” and “know” responses (Sharot et al., 2004; Wiesmann and Ishai, 2008). Altogether, our results align with the view that hippocampal structures, ventromedial prefrontal cortex, and posterior cingulate cortex form a circuit dedicated to the retrieval of personal memories including rich contextual information (Kensinger & Schacter, 2007; Maratos et al., 2001; Sterpenich et al., 2006). Interestingly, this network was also highly sensitive to the emotional content of scenes, as discussed below.

### 4.4 Emotional modulation of memory reactivation

In the FFA ROI reactivated during face retrieval, we observed a significant emotional increase for MA trials, but also to some degree CR, while there was no further enhancement during correct retrieval (Hit). Thus, during MA, negative scene cues evoked a high activation level in FFA, similar to Hit, whereas neutral cues produced a weaker response, similar to Miss or CR. In parallel with these visual effects, MA cued by negative scenes also elicited greater activation in fronto-parietal networks associated with memory retrieval and subjective “oldness” (see Fig. 6).

Altogether, these data indicate that emotional signals from the cues tended to produce brain activity patterns resembling Hit when participants made MA errors, whereas neutral cues produced activity patterns closer to Miss during the same errors, suggesting that MA may result from different mechanisms depending on negative vs neutral context. Earlier studies showed that negative affect can boost recollection and familiarity (Ochsner, 2000a; Rimmele, Davachi, Petrov, et al., 2011) but impair memory associations (Bisby & Burgess, 2017), so it is conceivable that the emotional scene content enhanced familiarity signals and triggered greater reactivation in FFA, eventually favoring the retrieval of an incorrect face. Stronger feelings of remembering may result from a modulation of sensory representations and stored associations through top-down influences from emotional systems, even when operating on incorrect representations. This effect may predominate with negative over neutral cues because of more diffuse spreading of associations in long-term memory stores (Foster & Sahakyan, 2011), or because of reduced integration with precise contextual information in emotional situations (Adolphs et al., 2005; Brewin et al., 2010).

These findings add to an earlier study (Hofstetter et al., 2012), where we found enhanced fusiform reactivation for the retrieval of faces previously paired with negative cues. In the latter study, only Hit responses were considered (errors were minimized by design), and we did not probe for correct face identity on every trial. Therefore, the Hit responses in this study could represent true recollection (as here) but also misattributions, where participants responded “face” (rather than “word” or “new”) without remembering the exact face identity. However, unlike our earlier study, here we did not find an emotional modulation of FFA during Hits. This might result from several differences in the current paradigm, including higher memory load, longer delay till retrieval, or ceiling effects in visual reactivation for Hits relative to other conditions.

Finally, comparing Hit vs MA revealed a network including VMPFC, ACC, PCC, amygdala, hippocampus, and parahippocampal cortex, typically implicated in autobiographical memory (Cabeza et al., 2004; Fink et al., 1996), which was active with both negative and neutral cues during Hits, but also with negative cues during MA. These activations parallel the pattern found in FFA. Thus, while this network activated for true recollection independent of cue emotionality, a subpart was also active during spurious recollection cued by negative scenes. These data suggest that different mechanisms may be at play during true and spurious retrieval cued by emotional scenes. Emotional cues may induce a feeling of remembering without true recollection (Rimmele, Davachi, Petrov, et al., 2011; Sharot & Phelps, 2004), by promoting stronger reactivation of sensory and other cortical areas within memory networks.

Accordingly, our functional connectivity analysis revealed greater coupling between left FFA and left hippocampus during Hits cued with the negative vs neutral scenes. This suggests stronger recruitment of the hippocampus necessary for correct face retrieval and retrieval of specific contextual associations in emotional conditions. This coupling between FFA and hippocampus did not occur during MA despite increased subjective familiarity, suggesting that functional connectivity between these areas contributes to “true” memory recollection.

A prominent model of intrusive memories in PTSD (Brewin et al., 2010) proposed that stressful situations might bias memories toward sensory-based representations without contextual elaboration. Sensory memories stored in visual and temporal structures might be reactivated involuntarily in a bottom-up fashion by corresponding cues and thus cause flashback, whereas retrieval of contextual information would require additional activation of fronto-parietal areas. Our results do not provide evidence for bottom-up (but rather top-down) reactivation of visual areas in response to emotional cues, but accord with enhanced cortical reactivation in emotional contexts. Bottom-up reactivation may only apply for truly stressful situations, unlike the scene pictures used here.

### 4.5 Conclusions

Our study provides novel evidence for reactivation of face-selective visual cortex during face retrieval during both correct and incorrect recall, extending current knowledge on memory functioning and memory errors. We show that reactivation of the FFA follows the subjective experience of remembering, rather than actual information seen at encoding. We also show that emotion can enhance memory reactivation when familiarity signals are evoked by the cue but true contextual information is absent. Overall, our findings highlight the role of content-specific cortical reactivation in memory errors, as well as its emotional modulation. Such emotion effects on memory errors have important implications for real life situations (e.g., eye witness) where retrieval may be distorted by emotional cues (Loftus, 2003).

## 5 Declaration of competing interests

No competing interests declared.

## 6 Authors’ contributions

CH: Project administration, Writing – review & editing, Writing – original draft, Visualization, Validation, Software, Methodology, Investigation, Formal analysis, Data curation, Conceptualization MR: Writing – review & editing, Writing – original draft, Visualization, Validation, Software, Methodology, Formal analysis, Data curation, Conceptualization

KI: Writing – review & editing, Formal analysis

AP: Writing – review & editing, Methodology, Investigation, Data curation

PV: Writing – review & editing, Supervision, Resources, Project administration, Methodology, Funding acquisition, Conceptualization

## 7 Data availability statement

All data necessary to verify, interpret and extend published research will be freely available through a dedicated repository: https://osf.io/2y5re/overview?view_only=2053495d2ec140c581ae9ff43f399be9.

## 8 Ethics statement

In the present study, all participants provided written informed consent and were financially compensated. The local Research Ethics approved the experimental protocol of the study.

## Acknowledgments

This work was supported by the Academic Society of Geneva (Fund Foremane) and by the National Center of Competence in Research (NCCR) Affective Sciences financed by the Swiss National Science Foundation and hosted by the University of Geneva (grant No 51NF40-104897). MR received funding from the European Union’s Framework Programme for Research and Innovation Horizon Europe (2021-2027) under the Marie Skłodowska-Curie Grant Agreement No. 101064781. This study was conducted in the Brain and Behavior Lab (BBL; University of Geneva, Switzerland) and benefited from support of the BBL technical staff.

## Notes

### Competing Interest Statement

The authors have declared no competing interest.

## References

Adolphs, R., Denburg, N. L., & Tranel, D. (2001). The amygdala’s role in long-term declarative memory for gist and detail. Behavioral Neuroscience, 115(5), 983–992. 10.1037//0735-7044.115.5.983

Adolphs, R., Gosselin, F., Buchanan, T. W., Tranel, D., Schyns, P., & Damasio, A. R. (2005). A mechanism for impaired fear recognition after amygdala damage. Nature, 433(7021), 68–72. 10.1038/nature03086

Antypa, D., Vuilleumier, P., & Rimmele, U. (2019). Suppressing but not intensifying emotion decreases arousal and subjective sense of recollection. Emotion, 19(6), 950–963. 10.1037/EMO0000493

Bisby, J. A., & Burgess, N. (2017). Differential effects of negative emotion on memory for items and associations, and their relationship to intrusive imagery. Current Opinion in Behavioral Sciences, 17, 124–132. 10.1016/j.cobeha.2017.07.012

Bone, M. B., Ahmad, F., & Buchsbaum, B. R. (2020). Feature-specific neural reactivation during episodic memory. Nature Communications, 11(1), 1–13. 10.1038/s41467-020-15763-2

Bosch, S. E., Jehee, J. F. M., Fernández, G., & Doeller, C. F. (2014). Reinstatement of associative memories in early visual cortex is signaled by the hippocampus. The Journal of Neuroscience : The Official Journal of the Society for Neuroscience, 34(22), 7493–7500. 10.1523/JNEUROSCI.0805-14.2014

Bower, G. H. (1996). Reactivating a reactivation theory of implicit memory. Consciousness and Cognition, 5(1–2), 27–72. 10.1006/ccog.1996.0004

Bradley, M., & Lang, P. J. (1994). MEASURING EMOTION : THE SELF-ASSESSMENT SEMANTIC DIFFERENTIAL MANIKIN AND THE. 25(I).

Brett, M., Anton, J. L., Valabregue, R., & Poline, J. B. (2002). Region of interest analysis using an SPM toolbox. NeuroImage, 16, 497. 10.1016/S1053-8119(02)90010-8

Brewin, C. R., Gregory, J. D., Lipton, M., & Burgess, N. (2010). Intrusive Images in Psychological Disorders: Characteristics, Neural Mechanisms, and Treatment Implications. Psychological Review, 117(1), 210–232. 10.1037/A0018113

Buckner, R. L., Goodman, J., Burock, M., Rotte, M., Koutstaal, W., Schacter, D., Rosen, B., & Dale, A. M. (1998). Functional-Anatomic Correlates of Object Priming in Humans Revealed by Rapid Presentation Event-Related fMRI. Neuron, 20(2), 285–296. 10.1016/S0896-6273(00)80456-0

Cabeza, R., Prince, S. E., Daselaar, S. M., Greenberg, D. L., Budde, M., Dolcos, F., LaBar, K. S., & Rubin, D. C. (2004). Brain Activity during Episodic Retrieval of Autobiographical and Laboratory Events: An fMRI Study using a Novel Photo Paradigm. Journal of Cognitive Neuroscience, 16(9), 1583–1594. 10.1162/0898929042568578

Cavanna, A. E., & Trimble, M. R. (2006). The precuneus: A review of its functional anatomy and behavioural correlates. Brain, 129(3), 564–583. 10.1093/brain/awl004

Chen, Y. Y., Areti, A., Yoshor, D., & Foster, B. L. (2024). Perception and Memory Reinstatement Engage Overlapping Face-Selective Regions within Human Ventral Temporal Cortex. Journal of Neuroscience, 44(22). 10.1523/JNEUROSCI.2180-23.2024

Chiu, Y. C., Dolcos, F., Gonsalves, B. D., & Cohen, N. J. (2013). On opposing effects of emotion on contextual or relational memory. Frontiers in Psychology, 4(MAR), 2–5. 10.3389/fpsyg.2013.00103

Danker, J. F., & Anderson, J. R. (2010). The ghosts of brain states past: Remembering reactivates the brain regions engaged during encoding. Psychological Bulletin, 136(1), 87. 10.1037/A0017937

Daselaar, S. M., Fleck, M. S., Dobbins, I. G., Madden, D. J., & Cabeza, R. (2006). Effects of Healthy Aging on Hippocampal and Rhinal Memory Functions: An Event-Related fMRI Study. Cerebral Cortex, 16(12), 1771–1782. 10.1093/CERCOR/BHJ112

Dolcos, F., LaBar, K. S., & Cabeza, R. (2004). Interaction between the amygdala and the medial temporal lobe memory system predicts better memory for emotional events. Neuron, 42, 855–863. 10.1016/S0896-6273(04)00289-2

Eldridge, L. L., Knowlton, B. J., Furmanski, C. S., Bookheimer, S. Y., & Engel, S. A. (2000). Remembering episodes: A selective role for the hippocampus during retrieval. Nature Neuroscience, 3(11), 1149–1152. 10.1038/80671

Epstein, R., & Kanwisher, N. (1998). A cortical representation the local visual environment. Nature, 392(6676), 598–601. 10.1038/33402

Favila, S. E., Lee, H., & Kuhl, B. A. (2020). Transforming the Concept of Memory Reactivation. Trends in Neurosciences, 43(12), 939. 10.1016/J.TINS.2020.09.006

Fink, G. R., Markowitsch, H. J., Reinkemeier, M., Bruckbauer, T., Kassler, J., & Heiss, W. D. (1996). Cerebral Representation of One’s Own Past: Neural Networks Involved in Autobiographical Memory. Journal of Neuroscience, 16(13), 4275–4282. 10.1523/JNEUROSCI.16-13-04275.1996

Flanagin, V. L., Klinkowski, S., Brodt, S., Graetsch, M., Roselli, C., Glasauer, S., & Gais, S. (2023). The precuneus as a central node in declarative memory retrieval. 1–10.

Foster, N. L., & Sahakyan, L. (2011). The role of forget-cue salience in list-method directed forgetting. Memory, 19(1), 110–117. 10.1080/09658211.2010.537665

Gershman, S. J., Schapiro, A. C., Hupbach, A., & Norman, K. A. (2013). Neural context reinstatement predicts memory misattribution. Journal of Neuroscience, 33(20), 8590–8595. 10.1523/JNEUROSCI.0096-13.2013

Grunwald, T., Pezer, N., Münte, T. F., Kurthen, M., Lehnertz, K., Van Roost, D., Fernández, G., Kutas, M., & Elger, C. E. (2003). Dissecting out conscious and unconscious memory (sub)processes within the human medial temporal lobe. NeuroImage, 20(SUPPL. 1), S139–S145. 10.1016/J.NEUROIMAGE.2003.09.004

Hebscher, M., Ibrahim, C., & Gilboa, A. (2020). Precuneus stimulation alters the neural dynamics of autobiographical memory retrieval. NeuroImage, 210(January), 116575. 10.1016/j.neuroimage.2020.116575

Henke, K., Mondadori, C. R. A., Treyer, V., Nitsch, R. M., Buck, A., & Hock, C. (2003). Nonconscious formation and reactivation of semantic associations by way of the medial temporal lobe. Neuropsychologia, 41(8), 863–876. 10.1016/S0028-3932(03)00035-6

Hofstetter, C., Achaibou, A., & Vuilleumier, P. (2012). NeuroImage Reactivation of visual cortex during memory retrieval : Content speci fi city and emotional modulation. NeuroImage, 60(3), 1734–1745. 10.1016/j.neuroimage.2012.01.110

Jaeger, A., Konkel, A., & Dobbins, I. G. (2013). Unexpected novelty and familiarity orienting responses in lateral parietal cortex during recognition judgment. Neuropsychologia, 51(6), 1061–1076. 10.1016/J.NEUROPSYCHOLOGIA.2013.02.018

Kahn, I., Davachi, L., & Wagner, A. D. (2004). Functional-Neuroanatomic Correlates of Recollection: Implications for Models of Recognition Memory. Journal of Neuroscience, 24(17), 4172–4180. 10.1523/JNEUROSCI.0624-04.2004

Kanwisher, N., & Yovel, G. (2006). The fusiform face area: A cortical region specialized for the perception of faces. In Philosophical Transactions of the Royal Society B: Biological Sciences (Vol. 361, Issue 1476, pp. 2109–2128). Royal Society. 10.1098/rstb.2006.1934

Kellen, D., Winiger, S., Dunn, J. C., & Singmann, H. (2021). Testing the Foundations of Signal Detection Theory in Recognition Memory. Psychological Review, 128(6), 1022–1050. 10.1037/REV0000288

Kensinger, E. A., Garoff-Eaton, R. J., & Schacter, D. L. (2007). How negative emotion enhances the visual specificity of a memory. Journal of Cognitive Neuroscience, 19, 1872–1887. 10.1162/jocn.2007.19.11.1872

Kensinger, E. A., & Schacter, D. L. (2006). Amygdala activity is associated with the successful encoding of item, but not source, information for positive and negative stimuli. The Journal of Neuroscience : The Official Journal of the Society for Neuroscience, 26(9), 2564–2570. 10.1523/JNEUROSCI.5241-05.2006

Kensinger, E. A., & Schacter, D. L. (2007). Remembering the specific visual details of presented objects: neuroimaging evidence for effects of emotion. Neuropsychologia, 45(13), 2951–2962. 10.1016/J.NEUROPSYCHOLOGIA.2007.05.024

Kraemer, P. M., Fontanesi, L., Spektor, M. S., & Gluth, S. (2021). Response time models separate single- and dual-process accounts of memory-based decisions. Psychonomic Bulletin and Review, 28(1), 304–323. 10.3758/s13423-020-01794-9

Kurkela, K. A., & Dennis, N. A. (2016). Event-related fMRI studies of false memory: An Activation Likelihood Estimation meta-analysis. Neuropsychologia, 81, 149–167. 10.1016/J.NEUROPSYCHOLOGIA.2015.12.006

LaBar, K. S., & Cabeza, R. (2006). Cognitive neuroscience of emotional memory. Nature Reviews. Neuroscience, 7(1), 54–64. 10.1038/NRN1825

Lang, P. . J. ., Bradley, M. . M. ., & Cuthbert, B. . N. . (2008). International affective picture system (IAPS): Affective ratings of pictures and instruction manual. In Technical Report A-8. 10.1016/j.epsr.2006.03.016

Lee, T. M. C., Au, R. K. C., Liu, H. L., Ting, K. H., Huang, C. M., & Chan, C. C. H. (2009). Are errors differentiable from deceptive responses when feigning memory impairment? An fMRI study. Brain and Cognition, 69(2), 406–412. 10.1016/J.BANDC.2008.09.002

Li, M., Huang, H., Guo, B., & Meng, M. (2022). Distinct response properties between the FFA to faces and the PPA to houses. Brain and Behavior, 12(8), 1–14. 10.1002/brb3.2706

Loftus, E. F. (2003). Make-Believe Memories. American Psychologist, 58(11), 867–873. 10.1037/0003-066X.58.11.867

Maldjian, J. A., Laurienti, P. J., Kraft, R. A., & Burdette, J. H. (2003). An automated method for neuroanatomic and cytoarchitectonic atlas-based interrogation of fMRI data sets. NeuroImage, 19(3), 1233–1239. 10.1016/S1053-8119(03)00169-1

Maratos, E. J., Dolan, R. J., Morris, J. S., Henson, R. N., & Rugg, M. D. (2001). Neural activity associated with episodic memory for emotional context. Neuropsychologia, 39, 910–920. 10.1016/S0028-3932(01)00025-2

Marchewka, A., Brechmann, A., Nowicka, A., Jednoróg, K., Scheich, H., & Grabowska, A. (2008). False recognition of emotional stimuli is lateralised in the brain: An fMRI study. Neurobiology of Learning and Memory, 90(1), 280–284. 10.1016/J.NLM.2008.01.012

Mather, M., & Knight, M. (2008). The emotional harbinger effect: poor context memory for cues that previously predicted something arousing. Emotion (Washington, D.C.), 8(6), 850–860. 10.1037/A0014087

McClelland, J. L., McNaughton, B. L., & O’Reilly, R. C. (1995). Why there are complementary learning systems in the hippocampus and neocortex: insights from the successes and failures of connectionist models of learning and memory. Psychological Review, 102(3), 419–457. 10.1037/0033-295X.102.3.419

Meaux, E., Sterpenich, V., & Vuilleumier, P. (2019). Emotional learning promotes perceptual predictions by remodeling stimulus representation in visual cortex. Scientific Reports 2019 9:1, 9(1), 1–14. 10.1038/s41598-019-52615-6

Minear, M., & Park, D. C. (2004). A lifespan database of adult facial stimuli. Behavior Research Methods, Instruments, & Computers 2004 36:4, 36(4), 630–633. 10.3758/BF03206543

Ochsner, K. N. (2000a). Are affective events richly recollected or simply familiar? The experience and process of recognizing feelings past. Journal of Experimental Psychology. General, 129(2), 242–261. 10.1037/0096-3445.129.2.242

Ochsner, K. N. (2000b). Are affective events richly recollected or simply familiar? The experience and process of recognizing feelings past. Journal of Experimental Psychology: General, 129(2), 242–261. 10.1037/0096-3445.129.2.242

Paller, K. A., & Voss, J. L. (2004). Memory reactivation and consolidation during sleep. Learning & Memory, 11(6), 664. 10.1101/LM.75704

Pierce, B. H., & Kensinger, E. A. (2011). Effects of emotion on associative recognition valence and retention interval matter. 11(1), 139–144. 10.1037/a0021287.Effects

Pine, A., Sadeh, N., Ben-Yakov, A., Dudai, Y., & Mendelsohn, A. (2018). Knowledge acquisition is governed by striatal prediction errors. Nature Communications, 9(1), 1–14. 10.1038/s41467-018-03992-5

Polyn, S. M. (2005). Category-Specific Cortical Activity Precedes Retrieval During Memory Search. Science, 310(5756), 1963–1966. 10.1126/science.1117645

Polyn, S. M., Natu, V. S., Cohen, J. D., & Norman, K. A. (2005). Neuroscience: Category-specific cortical activity precedes retrieval during memory search. Science, 310(5756), 1963–1966. 10.1126/SCIENCE.1117645/SUPPL_FILE/POLYN.SOM.PDF

Rimmele, U., Davachi, L., Petrov, R., Dougal, S., & Phelps, E. A. (2011). Emotion enhances the subjective feeling of remembering, despite lower accuracy for contextual details. Emotion, 11(3), 553. 10.1037/A0024246

Rimmele, U., Davachi, L., & Phelps, E. a. (2011). Emotion enhances the subjective feeling of remembering, despite lower accuracy for contextual details. Emotion, 11(3). 10.1037/a0024246.Emotion

Rissman, J., Greely, H. T., & Wagner, A. D. (2010). Detecting individual memories through the neural decoding of memory states and past experience. Proceedings of the National Academy of Sciences of the United States of America, 107(21), 9849–9854. 10.1073/PNAS.1001028107/SUPPL_FILE/PNAS.201001028SI.PDF

Ritchey, M., & Cooper, R. A. (2020). Deconstructing the Posterior Medial Episodic Network. Trends in Cognitive Sciences, 24(6), 451–465. 10.1016/j.tics.2020.03.006

Rolls, E. T., Joliot, M., & Tzourio-Mazoyer, N. (2015). Implementation of a new parcellation of the orbitofrontal cortex in the automated anatomical labeling atlas. NeuroImage, 122, 1–5. 10.1016/j.neuroimage.2015.07.075

Schacter, D. L., & Buckner, R. L. (1998). Priming and the Brain. Neuron, 20(2), 185–195. 10.1016/S0896-6273(00)80448-1

Sharot, T., & Phelps, E. A. (2004). How arousal modulates memory: Disentangling the effects of attention and retention. *Cognitive*, Affective & Behavioral Neuroscience, 4(3), 294–306. 10.3758/CABN.4.3.294

Sinclair, A. H., & Barense, M. D. (2019). Prediction Error and Memory Reactivation: How Incomplete Reminders Drive Reconsolidation. Trends in Neurosciences, 42(10), 727–739. 10.1016/J.TINS.2019.08.007

St. Jacques, P. L., Olm, C., & Schacter, D. L. (2013). Neural mechanisms of reactivation-Induced updating that enhance and distort memory. Proceedings of the National Academy of Sciences of the United States of America, 110(49), 19671–19678. 10.1073/PNAS.1319630110/SUPPL_FILE/PNAS.201319630SI.PDF

Staresina, B. P., & Wimber, M. (2019). A Neural Chronometry of Memory Recall. Trends in Cognitive Sciences, 23(12), 1071–1085. 10.1016/J.TICS.2019.09.011

Stark, C. E. L., Okado, Y., & Loftus, E. F. (2010). Imaging the reconstruction of true and false memories using sensory reactivation and the misinformation paradigms. Learning & Memory, 17(10), 485–488. 10.1101/LM.1845710

Sterpenich, V., D’argembeau, A., Desseilles, M., Balteau, E., Albouy, G., Vandewalle, G., Degueldre, C., Luxen, A., Collette, F., Maquet, P., Ve Albouy, G., Vandewalle, G., Degueldre, C., Luxen, A., Collette, F., & Maquet, P. (2006). The locus ceruleus is involved in the successful retrieval of emotional memories in humans. The Journal of Neuroscience : The Official Journal of the Society for Neuroscience, 26(28), 7416–7423. 10.1523/JNEUROSCI.1001-06.2006

Swets, J. A., Green, D. M., Getty, D. J., & Swets, J. B. (1978). Signal detection and identification at successive stages of observation. Perception & Psychophysics 1978 23:4, 23(4), 275–289. 10.3758/BF03199711

Talmi, D., Schimmack, U., Paterson, T., & Moscovitch, M. (2007). The role of attention and relatedness in emotionally enhanced memory. Emotion, 7(1), 89–102. 10.1037/1528-3542.7.1.89

Tulving, E., & Thomson, D. M. (1973). Encoding specificity and retrieval processes in episodic memory. Psychological Review, 80(5), 352–373. 10.1037/H0020071

Vaden, K. I., Gebregziabher, M., Kuchinsky, S. E., & Eckert, M. A. (2012). Multiple imputation of missing fMRI data in whole brain analysis. NeuroImage, 60(3), 1843–1855. 10.1016/J.NEUROIMAGE.2012.01.123

Vuilleumier, P. (2005). How brains beware: Neural mechanisms of emotional attention. Trends in Cognitive Sciences, 9(12), 585–594. 10.1016/j.tics.2005.10.011

Vuilleumier, P., Schwartz, S., Clarke, K., Husain, M., & Driver, J. (2002). Testing Memory for Unseen Visual Stimuli in Patients with Extinction and Spatial Neglect. Journal of Cognitive Neuroscience, 14(6), 875–886. 10.1162/089892902760191108

Weidemann, C. T., & Kahana, M. J. (2016). Assessing recognition memory using confidence ratings and response times. Royal Society Open Science, 3(4), 150670. 10.1098/rsos.150670

Wheeler, M. E., & Buckner, R. L. (2003). Functional Dissociation among Components of Remembering: Control, Perceived Oldness, and Content. Journal of Neuroscience, 23(9), 3869–3880. 10.1523/JNEUROSCI.23-09-03869.2003

Wickens, T. D. (2002). Elementary signal detection theory. In New York (Vol. 1). 10.1093/acprof:oso/9780195092509.001.0001

Windmann, S., & Kutas, M. (2001). Electrophysiological Correlates of Emotion-Induced Recognition Bias. Journal of Cognitive Neuroscience, 13(5), 577–592. 10.1162/089892901750363172

Xue, G. (2018). The Neural Representations Underlying Human Episodic Memory. In Trends in Cognitive Sciences (Vol. 22, Issue 6). 10.1016/j.tics.2018.03.004

Xue, G. (2022). From remembering to reconstruction: The transformative neural representation of episodic memory. Progress in Neurobiology, 219, 102351. 10.1016/J.PNEUROBIO.2022.102351

Yonelinas, A. P., & Ritchey, M. (2015). The slow forgetting of emotional episodic memories: an emotional binding account. Trends in Cognitive Sciences, 19(5), 259–267. 10.1016/j.tics.2015.02.009

Yu, J., Tao, Q., Zhang, R., Chan, C. C. H., & Lee, T. M. C. (2019). Can fMRI discriminate between deception and false memory? A meta-analytic comparison between deception and false memory studies. Neuroscience and Biobehavioral Reviews, 104, 43–55. 10.1016/J.NEUBIOREV.2019.06.027

