## Supplementary materials for "Emotional modulation of visual cortical reactivation during true and false memories"

Title:

### Supplementary materials

#### 1. RT analysis

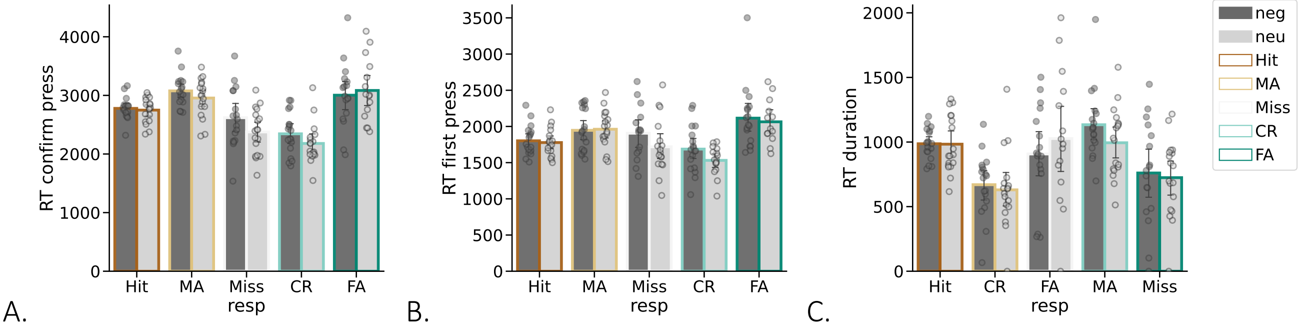


**Fig. S1** Reaction times (RT) on the scene-cued associative retrieval task as a function of memory response and emotion. **A.** RTs for the first button press - response choice, as a proxy of initial memory access. **B.** RTs for the second button press – response confirmation, as a proxy of memory decision. **C.** RT differences between the two button presses as a more precise timing of the memory decision after initial memory access.

#### 2. Functional localizer task results

Face-responsive areas were determined at the group level by identifying voxels differentially activated in the contrast “faces vs. [houses, words, patterns]”, masked inclusively [p = .005] by the anatomical region of fusiform gyrus from AAL2 (Rolls et al., 2015). For the resulting activation maps, a voxel-wise height threshold of $p < .001$ (uncorrected) was combined with a cluster level of $k = 10$ and used as a functionally-defined region of interest (ROI) for left ($k = 27$) and right ($k = 140$) fusiform face area (FFA).

As a control, place-responsive areas (Epstein & Kanwisher, 1998; Li et al., 2022) were determined at the group level by identifying voxels differentially activated in the contrast “houses vs. faces, words, patterns”, masked inclusively [p = .005] by the anatomical region of the parahippocampal cortex from the AAL2 (Rolls et al., 2015). For the resulting activation maps, a voxel-wise height threshold of $p < .001$ (uncorrected) was combined with a cluster level of $k = 10$ and used as a functionally defined region of interest (ROI) for left ($k = 49$) and right ($k = 83$) parahippocampal place area (PPA).

A.

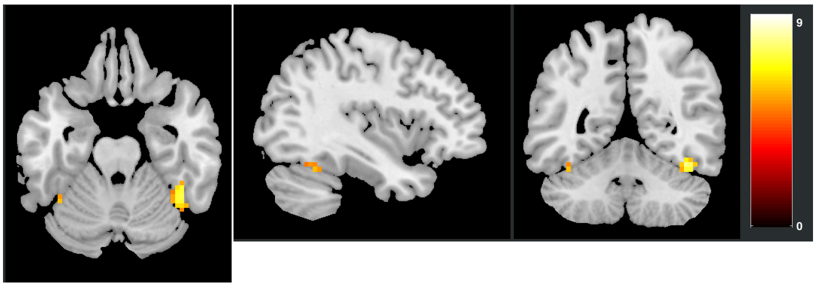

B.

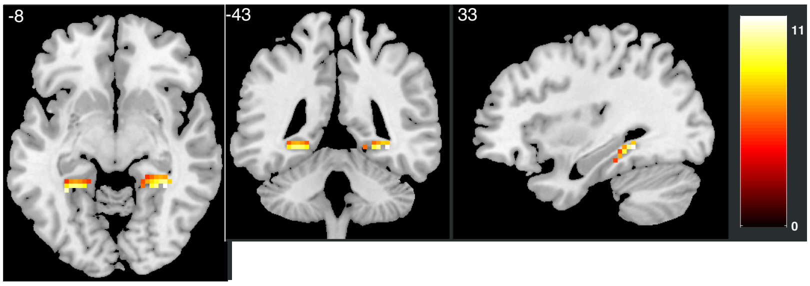


**Fig. S2** Results of the functional localizer task: A. Face-responsive areas (FFA), B. Place-responsive areas (PPA).

### 3. Analysis of control ROIs

Beta values extracted from the left PPA showed no modulation by emotion [$F(1,13) = 2.379$, $p = .147$, $\eta^2 = .007$], no modulation by memory response [$F(1,52) = 1.808$, $p = .188$, $\eta^2 = .026$], and no interaction between emotion and memory response [$F(1,52) = .320$, $p = .773$, $\eta^2 = .004$]. Beta values extracted from the right PPA showed only modulation by emotion [$F(1,13) = 5.602$, $p = .034$, $\eta^2 = .006$], with more activation for memories cued by negative scene but no effect of memory response [$F(1,52) = 1.895$, $p = .173$, $\eta^2 = .019$], and no interaction effect [$F(1,52) = .647$, $p = .566$, $\eta^2 = .004$].

For beta values extracted from the left amygdala, we observed no modulation by emotion [$F(1,13) = 2.247$, $p = .158$, $\eta^2 = .013$], and no interaction effect [$F(1,52) = 1.524$, $p = .239$, $\eta^2 = .039$]. However, we observed a modulation by memory response [$F(1,52) = 3.551$, $p = .020$, $\eta^2 = .795$], with activation to CR being higher than to MA [$t(13) = 3.334$, $p = .048$], and activation to Hits being higher than to MA [$t(13) = 3.847$, $p = .02$]. For beta values extracted from the right amygdala, we observed a modulation by emotion [$F(1,13) = 1.888$, $p = .006$, $\eta^2 = .029$], no modulation by memory response [$F(1,52) = 1.688$, $p = .203$, $\eta^2 = .04$] and no interaction effect [$F(1,52) = 1.644$, $p = .2$, $\eta^2 = .02$].

In the beta values extracted from the left hippocampus, we observed no modulation by emotion [$F(1,13) = 1.166$, $p = .3$, $\eta^2 = .006$], no modulation by memory response [$F(1,52) = 2.066$, $p = .099$, $\eta^2 = .053$], and no interaction between emotion and memory response [$F(1,52) = 1.06$, $p = .363$, $\eta^2 = .021$]. In the beta values extracted from the right hippocampus, we observed a trend of modulation by emotion [$F(1,13) = 4.03$, $p = .066$, $\eta^2 = .012$], no modulation by memory response [$F(1,52) = .98$, $p = .391$, $\eta^2 = .025$], and no interaction between emotion and memory response [$F(1,52) = .992$, $p = .398$, $\eta^2 = .011$].

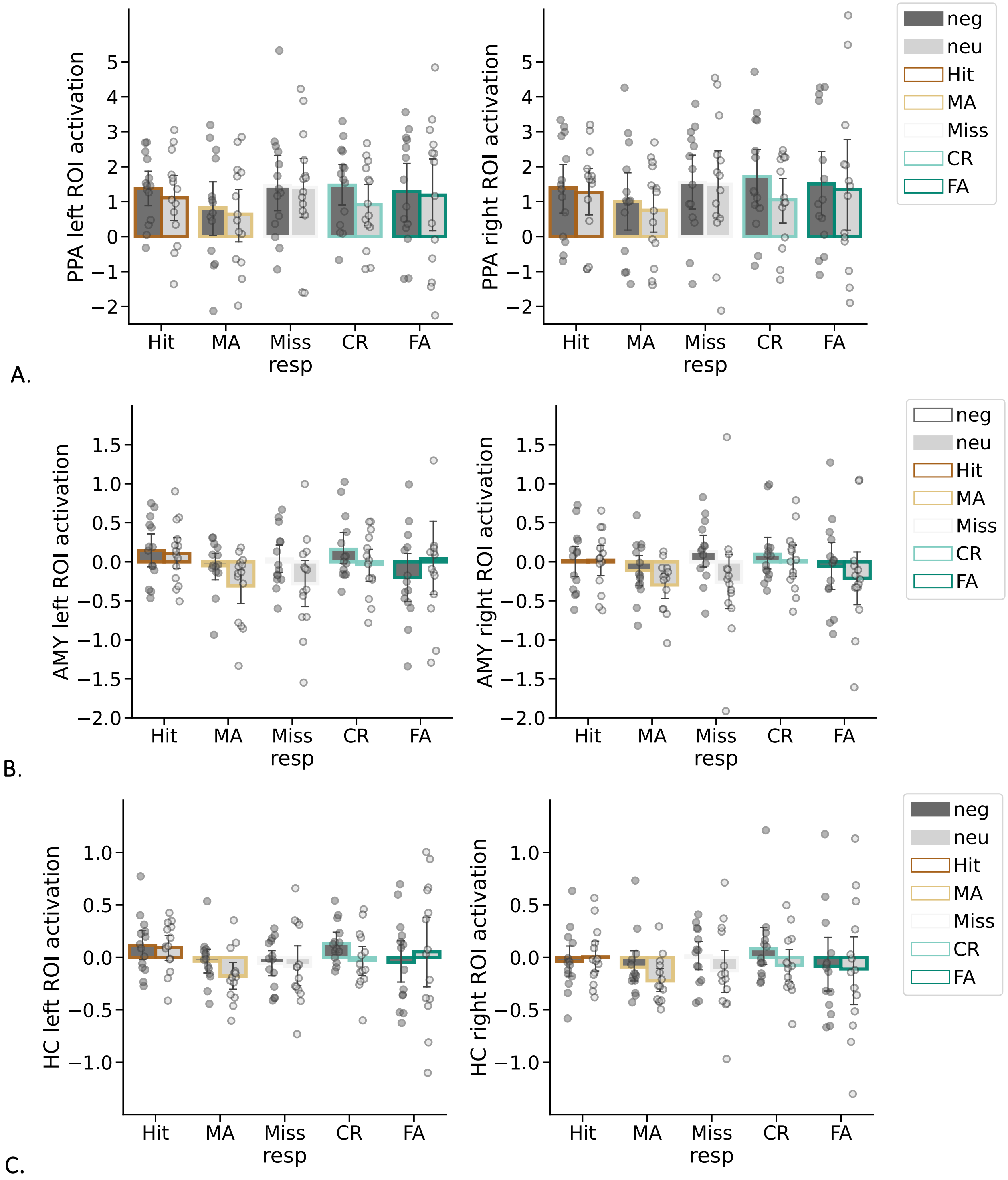


**Fig. S3** Modulation of control ROIs by memory condition and emotional context. **A**. Parahippocampal Place Area (PPA) left and right, defined in both hemispheres by an independent face localizer scan; **B**. Amygdala left and right; **C.** Hippocampus left and right.

**Table S1** Results of the Functional Localizer analysis. A voxel-wise height threshold of p < .001 (uncorrected), combined with a cluster level of k = 1.

Table shows all local maxima separated by more than 8mm. Regions were automatically labelled using the AAL-3 atlas, x, y, and z Montreal Neurological Institute (MNI) coordinates in the lef-right, anterior-posterior, and inferior-superior dimensions, respectively, pFWE signifies p-values at the peak level.
